# Astrocytic extracellular vesicles modulate neuronal calcium homeostasis via transglutaminase-2

**DOI:** 10.1101/2021.09.30.462507

**Authors:** Elisa Tonoli, Ivan Verduci, Ilaria Prada, Martina Gabrielli, Greta Forcaia, Clare Coveney, Maria Pia Savoca, David J. Boocock, Giulio Sancini, Michele Mazzanti, Claudia Verderio, Elisabetta A.M. Verderio

## Abstract

We have uncovered a novel role for astrocytes-derived extracellular vesicles (EVs) in controlling intraneuronal Ca^2+^ concentration ([Ca^2+^]_i_) and identified transglutaminase-2 (TG2) as a surface-cargo of astrocytes-derived EVs. Incubation of hippocampal neurons with primed astrocyte-derived EVs have led to an increase in [Ca^2+^]_i_, unlike EVs from TG2-knockout astrocytes. Exposure of neurons or brain slices to extracellular TG2 promoted a [Ca^2+^]_i_ rise, which was reversible upon TG2 removal and was dependent on Ca^2+^ influx through the plasma membrane. Patch-clamp and calcium imaging recordings revealed TG2-dependent neuronal membrane depolarisation and activation of inward currents, due to the opening of L-type-VOCCs and to Na^+^/Ca^2+^ -exchanger (NCX) operation in the reverse mode, as indicated by VOCCs/NCX pharmacological inhibitors. A subunit of Na^+^/K^+^-ATPase was selected by comparative proteomics and identified as being functionally inhibited by extracellular TG2, implicating Na^+^/K^+^-ATPase inhibition in NCX reverse mode-switching leading to Ca^2+^ influx and higher basal [Ca^2+^]_i_. These data suggest that reactive astrocytes control intraneuronal [Ca^2+^]_i_ through release of EVs with TG2 as responsible cargo, which could have a significant impact on synaptic activity in brain inflammation.

## Introduction

The regulation of Ca^2+^ is a fundamental biological process in neurons, with Ca^2+^ signalling as the basis of synaptic plasticity, neurons survival and their ability to communicate (Catterall & Few, 2008; Marambaud, Dreses-Werringloer et al., 2009). In neurons basal Ca^2+^ levels are tightly regulated within a narrow physiological range (Carafoli, 1987; Jones & Keep, 1988), and a rise in intracellular Ca^2+^ concentration ([Ca^2+^]_i_) is the major trigger of neurotransmitter release from nerve endings. Propagation of action potentials and membrane depolarisation (∼ 20 mV) produces Ca^2+^ influx through voltage-dependent Ca^2+^ channels which reaches micromolar levels at highly localised presynaptic sites in axons (Forti, Pouzat et al., 2000). Since the smallest change in Ca^2+^ currents can have a dramatic impact on neuronal function, Ca^2+^ influx is controlled by the action of Ca^2+^ membrane transporters at the plasma membrane, such as the Ca^2+^ pump ATPase and the sodium/calcium (Na^+^/Ca^2+^)-exchanger (NCX), and it is buffered by a large set of Ca^2+^ binding proteins which act as modulators of cellular Ca^2+^ transients. Extensive literature indicates that Ca^2+^ dyshomeostasis in neurons is linked with aging and neurodegeneration. However, mechanism(s) leading to [Ca^2+^]_i_ alterations are still not completely clear (Nikoletopoulou & Tavernarakis, 2012; Oh, Oliveira et al., 2013).

Transglutaminase-2 (TG2) is a Ca^2+^-dependent crosslinking enzyme known to be involved in multiple neurodegenerative diseases linked to Ca^2+^ dysregulation (Grosso & Mouradian, 2012), as well as a variety of conditions triggered by inflammatory processes, such as wound-healing and tissue fibrosis (Verderio, Furini et al., 2015). The implication of TG2 was first suggested by the observation that the enzyme, in the presence of activating [Ca^2+^]_i_, is able to crosslink pathogenic misfolded proteins typical of neurodegenerative conditions (e.g. amyloid-β and α-synuclein), thus favouring formation of protein aggregates (Andringa, Lam et al., 2004; Benilova, Karran et al., 2012; Grosso, Woo et al., 2014; Hartley, Zhao et al., 2008; Junn, Ronchetti et al., 2003). However a number of recent observations, such as the potentiation of Ca^2+^-induced hippocampal damage by TG2 in mice brain and higher sensitivity to kainic acid-induced seizures (Tucholski, Roth et al., 2006), as well as the neurotoxic role of astrocytic TG2 following acute brain injury (Feola, Barton et al., 2017; Monteagudo, Feola et al., 2018), have hinted to a possible role of TG2 in excitotoxicity-induced neuronal cell death, which may be either consequent to [Ca^2+^]_i_ increases or, as a new hypothesis, triggered by TG2-mediated changes in [Ca^2+^]_i_. Astrocytes are an abundant source of TG2, which is externalised and accumulates in the extracellular matrix (ECM) in response to inflammatory stimuli (Pinzon, Breve et al., 2017). Importantly, astrocytes are key mediators of brain immune responses (Colombo & Farina, 2016) and TG2 has been shown to play a role in neuroinflammation (Ientile, Curro et al., 2015). Astrocytes control synapses either by direct contact and/or by secreted factors, released as single molecules or packaged into extracellular vesicles (EVs), which target pre- and post-synapses thus regulating synaptic behaviour (Farhy-Tselnicker & Allen, 2018). In this study we have hypothesised that extracellular TG2 could disrupt Ca^2+^ homeostasis in neurons. We have uncovered a new role for extra-neuronal TG2, released by reactive astrocytes via EVs, in increasing neuronal [Ca^2+^]_i_, and shown that this occurs through the opening of L-type VOCCs and NCX operation in the reverse mode, via regulation of Na^+^/K^+^-ATPase activity. This process is a novel example of neuron-regulation by astrocytes-derived EVs cargo under both physiological and neuroinflammatory conditions.

## Results

### TG2 released by primary astrocytes in association with EVs increases [Ca^2+^]_i_ in neurons

Previous studies have shown that astrocytes are a main source of extracellular TG2 in the brain, particularly in situations of inflammation/astroglial response to injury (Colak & Johnson, 2012; Monteagudo et al., 2018; Pinzon et al., 2017; Quinn, Yunes-Medina et al., 2018). A granular pattern of TG2 distribution was detected by immunofluorescence in either permeabilized or non-permeabilized fixed cultures of astrocytes, suggesting that TG2 has a predominant extracellular location in astrocytes (Fig. 1A). Evaluation of endogenous cell surface TG2 by an optimised activity assay revealed high extracellular TG2 activity (sensitive to the specific inhibitor ZDON) in cultured astrocytes (Fig. EV1). Conversely in hippocampal neurons, TG2 showed a significantly higher staining in permeabilized cells (Fig.1B), where it partially localised at synaptic sites, as indicated by co-staining with the pre- and post-synaptic markers VGLUT1 (blue) and SHANK2 (red) (Fig. 1C). Western blotting analysis of crude synaptosomes from adult mouse brain, revealed a significant enrichment of TG2 at synaptic contacts (Fig. 1D), confirming the presence of TG2 at synaptic sites *in vivo*, either in neuron terminals (validated by the expression of NR2B, PSD95, VGAT and VGLUT1) or in the astrocytic processes wrapping the synapses (validated by GFAP expression) (Fig. 1A and 1D) (Gylys, Fein et al., 2000).

**Figure 1.**
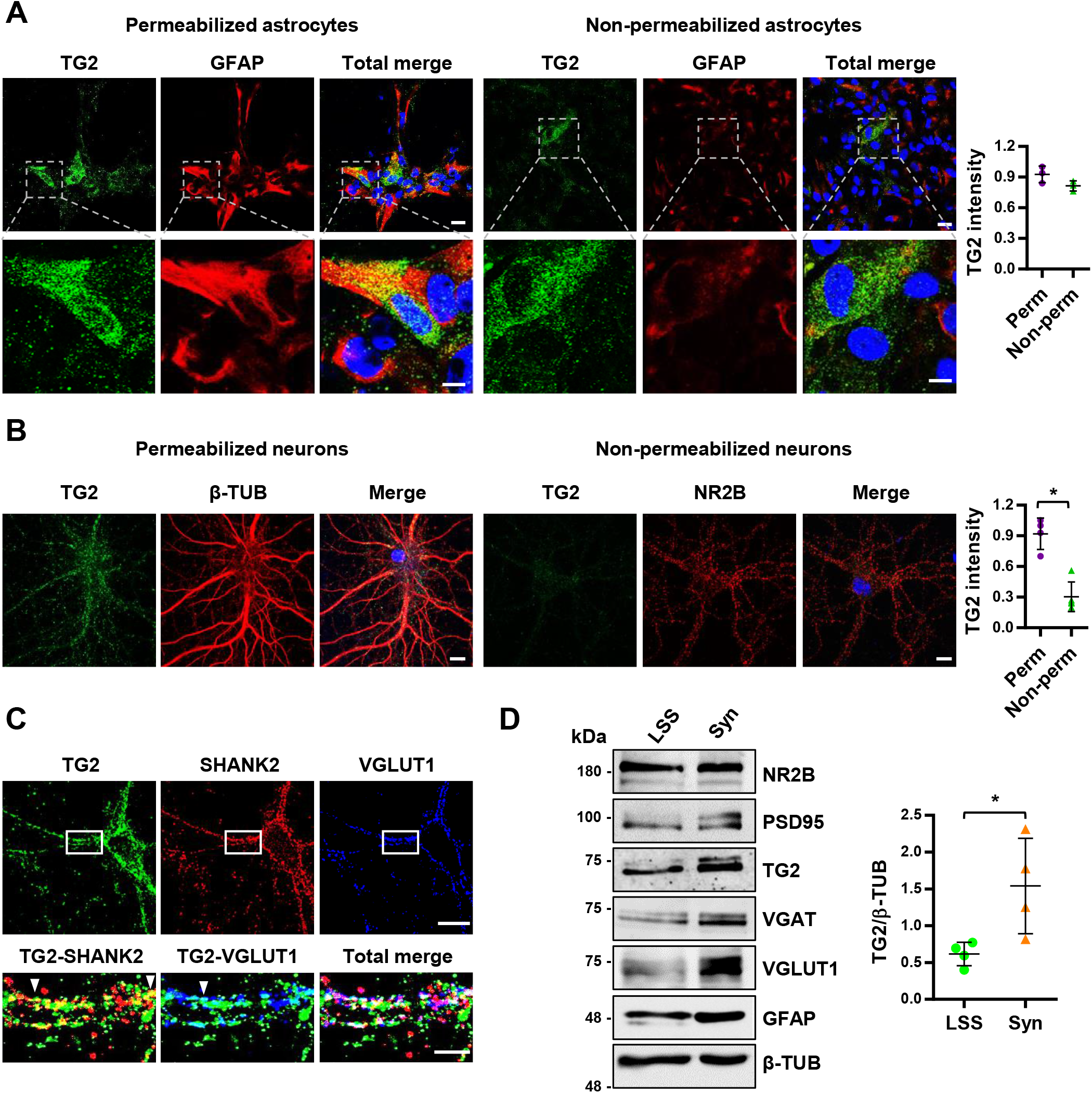
TG2 localises extracellularly in primary astrocytes and at synaptic sites in neurons. **A)** Immunofluorescence staining of primary astrocytes. Cells were fixed in 4% paraformaldehyde - 4 % sucrose (w/v), permeabilized (left panels) or non-permeabilized (right panel) and stained with anti-TG2 IA12 (green), DAPI (blue) and astrocytic marker anti-GFAP (red). Coverslips were visualised by laser scanning Leica SP5 confocal microscope using 63X oil immersion objective. Successive serial optical sections (1 µm) were recorded over 8 µm planes. Scale bar 20 µm. TG2 intensity was calculated by Leica software, divided by number of nuclei, and normalised to permeabilized values. Data is expressed as mean ± SD (N=3, Mann-Whitney test: p=NS). **B)** Neurons at 12 DIV were fixed and permeabilized (left panel) or non-permeabilized (right panel) and stained with anti-TG2 IA12 (green), anti-β-TUB or anti-NR2B (red) antibodies and DAPI (blue). Scale bar 10 µm. TG2 intensity was calculated as described in A (N=3, Mann-Whitney test: *p<0.05). **C)** Neurons at 22 Days In Vitro (DIV) were fixed, permeabilized and stained with anti-TG2 IA12 (green), anti-SHANK2 (red) and anti-VGLUT1 (blue) antibodies (N=3). Coverslips were visualised as described in A. Scale bar 20 µm (low magnification) and 5 µm (high magnification). **D)** Crude synaptosomes were isolated from adult mouse brain and probed with pre- and post-synaptic markers and for astrocyte marker GFAP. B-tubulin was used as loading control. TG2 is enriched in the synaptic fraction (Syn) compared to Low Speed Supernatant (LSS) representing the total lysate. Data is expressed as mean ± SD (N=4 independent experiments, Mann-Whitney test: *p<0.05).

Once externalised at the astrocyte surface TG2 can bind to matrix fibronectin (FN) (Pinzon et al., 2017) and our data confirmed a partial co-localisation of TG2 with FN (Fig. EV2). We hypothesised that TG2 may be released via extracellular vesicles (EVs), as recently described in other cell systems (Antonyak, Li et al., 2011; Diaz-Hidalgo, Altuntas et al., 2016; Furini, Schroeder et al., 2018). In this form, TG2 could travel to neurons, unless released from astrocytes in direct contact with synapses. To test this possibility, astrocyte-derived EVs were isolated from conditioned medium of primary astrocytes obtained from wild type (TGM2^+/+^) and knockout (TGM2^-/-^) mouse brain (De Laurenzi & Melino, 2001). Characterisation by Nanoparticle tracking analysis (Fig. 2A), TEM (Fig. 2B), detection of vesicular markers ALIX and Flotillin-2 and absence of negative marker TOM20 confirmed the size, morphology and vesicular cargo of EVs (Fig. 2C). NTA showed a trend increase in EVs number upon LPS treatment (Fig. 2A). Neither average particle number nor mode size were significantly different between WT and TG2KO cells, either untreated or LPS-treated (Fig. 2A-I and 2A-II), whereas a significant difference was observed between WT and WT+LPS particle number when looking at specific sizes (45-195 nm range) (Fig. 2A-III). TEM analysis and immunogold labelling of WT vesicles showed that they had the expected cup shape and confirmed the presence on the EVs surface of specific EVs marker CD63 and of TG2 (Fig. 2B). Western blotting analysis of astrocytes-derived EVs lysates revealed the presence of TG2 protein in WT EVs which displayed a trend increase upon pro-inflammatory stimulation with LPS (Fig. 2C). Furthermore, negligible TG2 was detected in vesicle-free supernatants after recovering all proteins by TCA precipitation, suggesting a prominent vesicular localisation of TG2 in primary astrocytes (Fig. 1A). These data show for the first time that TG2 is released from cultured astrocytes as a cargo of EVs under both resting physiological and stimulated (inflammatory) conditions. The localisation of TG2 in EVs was investigated using a sensitive enzymatic assay as we previously described (Furini et al., 2018). TG2 was detected at comparable levels in both intact and lysed EVs suggesting a predominant EVs surface location (Fig. 2D). To investigate the involvement of the astrocytes EVs TG2 cargo on neuronal function, the modulation of [Ca^2+^]_i_ by EVs isolated from WT and TG2KO LPS-primed astrocytes was monitored in fura-2-loaded hippocampal neurons. In order to exclude variations in [Ca^2+^]_i_ due to vesicular ATP leaking from astrocyte-derived EVs (D’Arrigo, Gabrielli et al., 2021), the assay was performed in the presence of the ATP degrading enzyme apyrase. As shown in Fig. 2E, TG2-containing EVs (EVs WT) were able to rise basal [Ca^2+^]_i_, whereas addition of TG2KO EVs did not cause significant changes. These data have two novel implications: that astrocytes-derived EVs concur to the regulation of neuronal Ca^2+^ homeostasis and that TG2, which is exposed on the surface of astrocytes-derived EVs, is responsible for this specific function.

**Figure 2.**
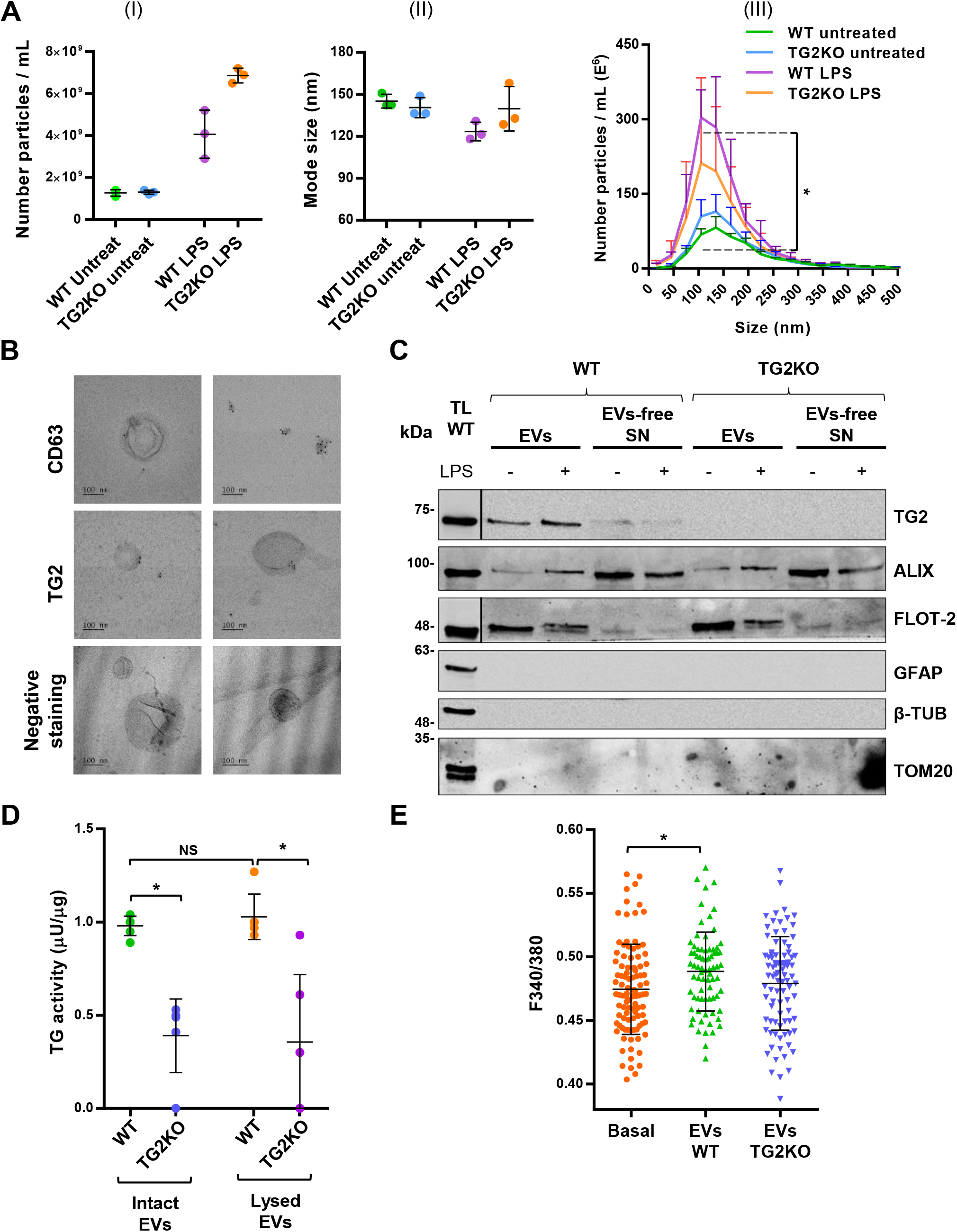
Astrocytic TG2 is released in association with EVs and modulates calcium homeostasis in neurons. **A)** EVs characterisation by nanoparticle tracking analysis (NTA). Average particle concentration (I) and mode size (II) of EVs isolated from untreated and LPS-treated WT and TG2KO astrocytes characterised by NTA (ZetaView, Particle Metrix). Data is shown as mean ± SD (N=3 independent experiments, Kruskal-Wallis Dunn’s test: p=NS). (III) Graph showing average particle concentration according to EVs size (Kruskal-Wallis Dunn’s test WT untreated – WT LPS: *p<0.05). **B)** EVs characterisation by Transmission electron microscopy (TEM). TEM analysis of EVs isolated from untreated WT astrocytes and stained with either anti-CD63 antibody (Abcam) or anti-TG2 antibody (IA12) and 6 nm gold colloidal anti-mouse IgG (top panels), or negative stained with uranyl acetate methylcellulose (bottom panel). **C)** Western blotting analysis of astrocytes-derived EVs and proteins TCA-precipitated from vesicle-free supernatants (EV-free SN) for the detection of EVs markers ALIX and FLOT-2, EVs negative marker TOM20, glial marker GFAP and of TG2. β-tubulin was used as loading control for TL. A representative blot of 3 independent experiments is shown. Treated cells were primed with LPS (1 µg/ml) for 24 h in serum-free medium before EVs isolation. Vertical black lines separate sections of the same membrane developed for different exposition times. **D)** Comparison of TG2 activity in intact and lysed EVs isolated from WT and TG2KO astrocytes. EVs were either resuspended as intact in PBS or lysed in a mild lysis buffer, to detect surface TG2 activity or total activity respectively. Data are expressed as mean ± SD normalised to WT intact EVs values (N=3 independent experiments; Kruskal-Wallis Dunn’s test: *p<0.05). **E)** Quantification of [Ca^2+^]_i_, expressed as F340/380, in 13 DIV neurons before and after stimulation with EVs isolated from LPS-treated WT or TG2KO astrocytes (N≥75 neurons from a preparation derived from ≥10 rat E18 embryos). EVs from either WT or TG2KO cells were resuspended in KRH supplemented with 30 U/ml of apyrase and added to neurons in the presence of 1 µM TTX, 50 μM APV and 10 μM CNQX. [Ca^2+^]_i_ was measured in multiple fields before (basal) and 20-60 min after addition of either WT or TG2KO EVs. Data are expressed as mean ± SD (one-way ANOVA test: *p<0.05).

### Extracellular TG2 increases cytoplasmic calcium concentration, both in cultured neurons and hippocampal slices

To gain insights into the mechanism leading to increased neuronal [Ca^2+^]_i_ by extracellular TG2 we next monitored neuronal [Ca^2+^]_i_ dynamics upon addition to the extracellular solution of purified TG2. Differentiated hippocampal neurons often exhibit synchronous Ca^2+^ oscillations in the soma, reflecting burst of neuronal firing (Bacci, Verderio et al., 1999). Incubation with purified TG2 promoted the onset of a synchronous Ca^2+^ spike and increased the interspike [Ca^2+^]_i_, leading to blockage of oscillatory activity (Fig. 3A-I, “TG2”). Upon TG2 removal by wash with KRH (Fig. 3A-I, “Wash”), interspike [Ca^2+^]_i_ decreased to pre-treatment values and spontaneous Ca^2+^ oscillations re-started, indicating that the action of soluble TG2 was reversible. To further examine the action of TG2 on [Ca^2+^]_i_, neurons were exposed to TG2 in the presence of TTX (1 µM), which prevents spontaneous synaptic and synchronous Ca^2+^ activity. In virtually all neurons tested, addition of TG2 led to a small but highly significant increase in [Ca^2+^]_i_ in the neuron cell bodies (Fig. 3A-II, “TG2”), while no [Ca^2+^]_i_ changes were observed in astrocytes present in the cultures. To rule out that the phenomenon was only found in dissociated cultures, we investigated the effect of TG2 on hippocampal slices. Similar Ca^2+^ response was evoked by TG2 in neurons in mouse hippocampal slices loaded with fura-2 when the TG2 protein was added in perfusion mode (Fig. 3B). In order to clarify whether extracellular TG2 increases basal Ca^2+^ levels by enhancing Ca^2+^ influx into neurons, we exposed cultured neurons to purified TG2 in Ca^2+^-free solution (Fig. 3C-I). No rise in basal [Ca^2+^]_i_ occurred in Ca^2+^-free conditions, while TG2-dependent Ca^2+^ rises were clearly recorded from the same neurons after addition of Ca^2+^ ions (2mM Ca^2+^) to the extracellular medium (Fig. 3C-II). This establishes that the [Ca^2+^]_i_ rise evoked by exogenous TG2 is mediated by Ca^2+^ influx from the extracellular environment through the plasma membrane. Interestingly, the action of TG2 was reversible, as observed in the absence of TTX (Fig. 3A-I) and neurons recovered basal [Ca^2+^]_i_ upon removal of the protein (“Wash”) from the extracellular solution (Fig. 3A-I, 3A-II and 3B). This finding implies that the observed process might not be due to transamidation, which leads to a covalent irreversible modification of protein substrates and should not disappear once TG2 is removed. To analyse this further, we tested the activity of exogenous TG2 added to hippocampal neurons during calcium imaging recordings and found that it was catalytically inactive in the same experimental conditions of the calcium imaging, unless activated by supplementation of a reducing agent (Fig. EV3). Notably, we previously showed that extracellular TG2 bound to the ECM is catalytically inactive or poorly active in the absence of a reducing agent (Verderio, Telci et al., 2003).

**Figure 3.**
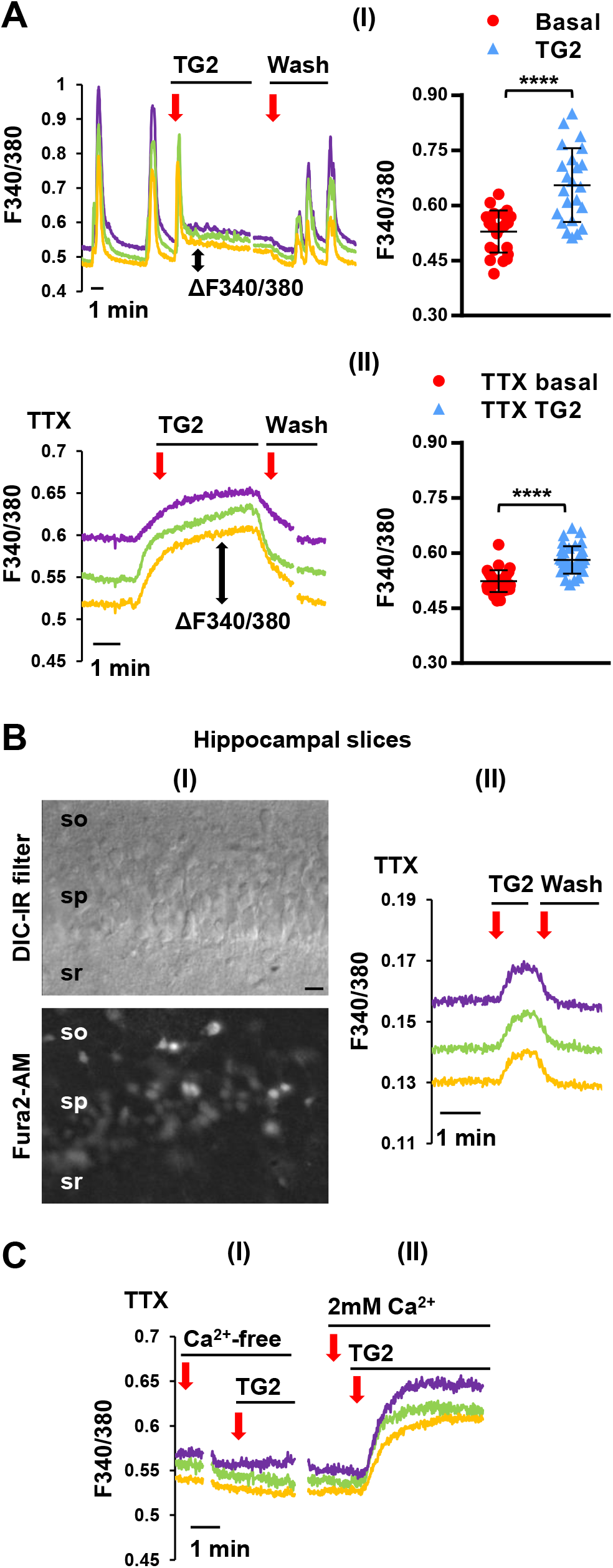
Exogenous TG2 increases interspike and basal [Ca^2+^]_i_ in hippocampal neurons in culture and in brain slices. **A)** (I) Representative temporal analysis of synchronous Ca^2+^ oscillations, expressed as F340/380, spontaneously occurring in 14 DIV Fura-2-loaded hippocampal neurons. Addition of exogenous TG2 promoted a Ca^2+^ spike followed by a sustained Ca^2+^ plateau (indicated by ΔF340/380), with a block of oscillations. Upon TG2 removal (wash with KRH) synchronous Ca^2+^ oscillations re-started. Three representative traces of neuronal [Ca^2+^]_i_ are shown in each graph. The graph shows the quantification of TG2-induced interspike [Ca^2+^]_i_ elevations compared with basal [Ca^2+^]_i_ (N=23 neurons, Student’s t-test: ****p<0.0001). (II) Addition of exogenous TG2 in the presence of TTX (1 µM) significantly increased [Ca^2+^]_i_ (indicated by ΔF340/380). Graph shows the quantification of TG2-induced [Ca^2+^]_i_ rises (peak Ca^2+^ rise) compared with basal [Ca^2+^]_i_ in the presence of TTX (N=33 neurons, Student’s t-test: ****p<0.0001). Data are presented as mean F340/380 ± SD. **B)** (I) Representative DIC (top) and 380 nm fluorescence (bottom) images of neurons in the stratum pyramidale (sp) of mouse hippocampal slices loaded with Fura-2/AM for calcium imaging experiments. The stratum oriens (so) and stratum radiatum (sr) are indicated above and below sp respectively. (II) Representative temporal plot of [Ca^2+^]_i_ rises evoked by perfusion of extracellular TG2 in fura-2 loaded hippocampal slices in 1 µM TTX (N=3 brain slices, 28 neurons in total). **C)** Temporal analysis of [Ca^2+^]_i_ changes in 14-20 DIV neurons exposed to exogenous TG2 in the absence (I) or in the presence (II) of extracellular Ca^2+^. Addition of TG2 in the absence of extracellular Ca^2+^ did not lead to changes in [Ca^2+^]_i_ (N=31 neurons).

We next asked whether TG2 secreted/externalised from neurons could rise basal [Ca^2+^]_i_ similarly to exogenous/astrocyte-derived TG2. To this aim, we first transfected hippocampal neurons with TG2 and measured [Ca^2+^]_i_. Transfected cells (pEGFP-N1-TG2) displayed an increase in resting [Ca^2+^]_i_ compared to control neurons (pEGFP-N1) (Fig. 4A). This suggests that upon over-expression neuronal TG2 may act extracellularly in an autocrine manner, controlling basal [Ca^2+^]_i_, albeit we cannot exclude an intracellular action of the protein. Treatment of the transfected cells with BOCDON, a non-permeable inhibitor of TG2 targeting extracellular TG2, did not affect changes in [Ca^2+^]_i_ evoked by the transfected TG2 (Fig. 4B). As testing the transamidating activity of TG2 solubilised in the extracellular saline confirmed that extracellular TG2 is predominantly inactive unless treated with a reducing agent (Fig. EV3), and that inhibition of extracellular TG2 does not affect intracellular Ca^2+^ changes, we conclude that TG2 evoking Ca^2+^ responses is prevalently inactive as a protein crosslinker.

**Figure 4.**
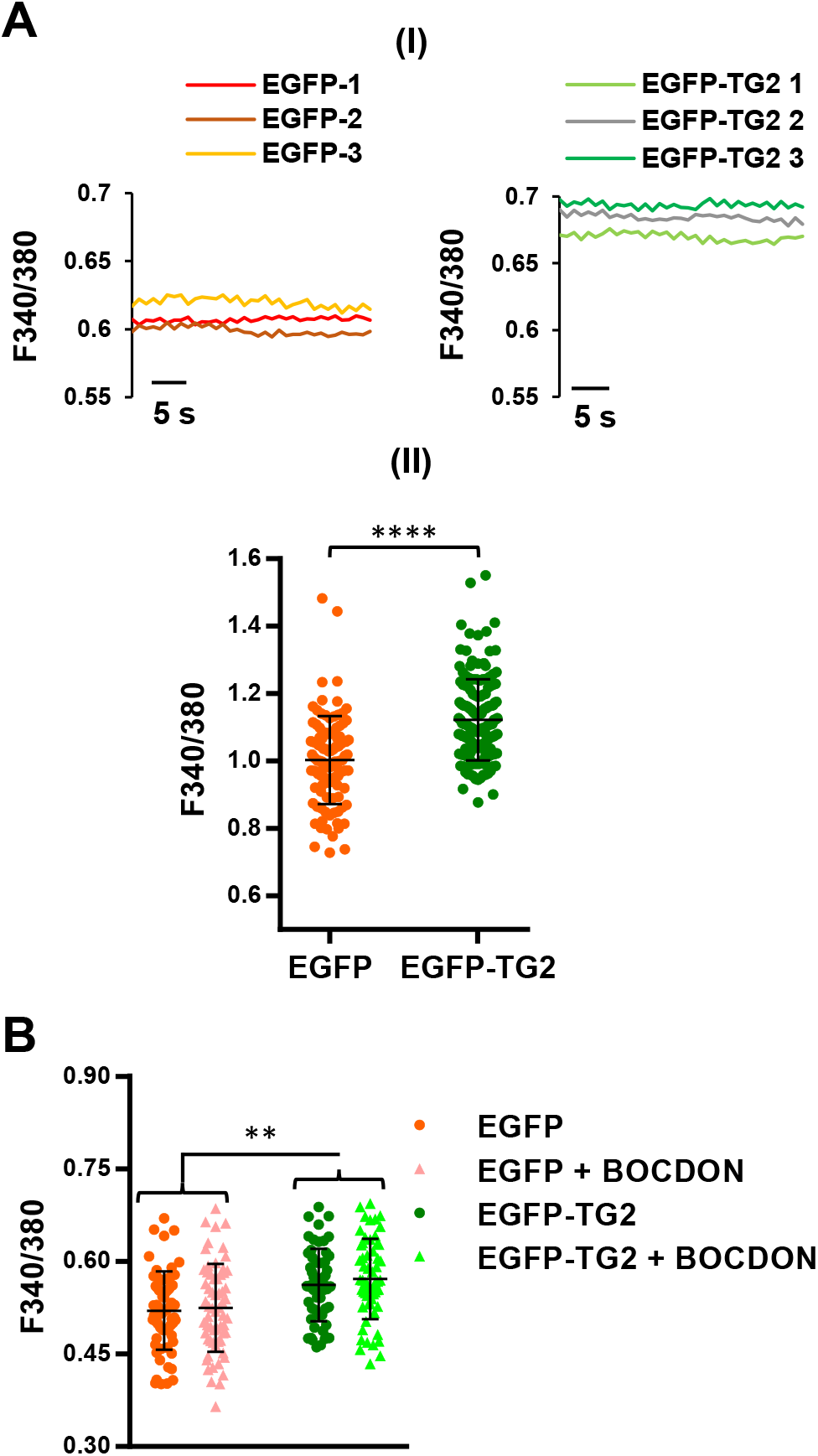
Transfected TG2 induces a calcium rise in an autocrine manner in neurons. **A)** (I) Representative traces showing basal [Ca^2+^]_i_, expressed as F340/380, in pEGFP-N1-TG2 or pEGFP-N1 transfected neurons (8 DIV). (II) Graph showing [Ca^2+^]_i_, in transfected neurons expressed as mean F340/380 ± SD normalised to EGFP (N>100 neurons. Mann-Whitney test: ****p<0.0001). **B)** A subgroup of transfected neurons was treated with the non-permeable TG2 inhibitor BOCDON (200 μM), which did not affect [Ca^2+^]_i_ (N>45 neurons). A significant difference in [Ca^2+^]_i_ between EGFP and EGFP-TG2 was observed in this subgroup as in graph A-II (one-way ANOVA test: **p<0.01).

### Extracellular TG2 causes calcium influx through L-type VOCCs

We next asked how extracellular TG2 promotes Ca^2+^ influx through the plasma membrane. Neurons were stimulated with exogenous TG2 in the presence of blockers of the main Ca^2+^ entry pathways in neurons, namely glutamate receptors and VOCCs. The NMDA receptor antagonist APV and the AMPA/kainate receptor antagonist CNQX did not affect Ca^2+^ increase upon TG2 addition, ruling out the involvement of these ligand-gated Ca^2+^ permeable receptors in the process (Fig. 5A-I and 5A-II). To analyse the possible contribution of VOCCs, we used a selection of pharmacological inhibitors of these channels. Pre-treatment with cadmium, a general blocker of VOCCs, reduced TG2-dependent Ca^2+^ responses by about 82.4% (Fig. 5B-I) and caused an almost complete recovery (88.3%) of [Ca^2+^]_i_ towards basal levels when applied during the plateau phase of Ca^2+^ response induced by exogenous TG2 (Fig. 5C-I). Similarly, pre-treatment with nickel, a more specific inhibitor of T-type VOCCs, decreased TG2-dependent Ca^2+^ rises by 66.2% (Fig. 5B-II) while it caused a drop of [Ca^2+^]_i_ below basal level (151% inhibition) when applied during the plateau phase (Fig. 5C-II). The involvement of VOCCs was further indicated by the observation that Ca^2+^ transients evoked by depolarisation (15 mM KCl) rose faster and reached a higher peak in the presence of extracellular TG2, with a significant increase in average Ca^2+^ influx (Fig. 5D). Among VOCCs controlling Ca^2+^ transport through the plasma membrane, L-type VOCCs are highly abundant in the somatodendritic region of hippocampal neurons (Condliffe, Corradini et al., 2010; Leitch, Szostek et al., 2009; Pravettoni, Bacci et al., 2000). We therefore asked whether these channels might contribute to Ca^2+^ influx in the neuronal soma evoked by exogenous TG2. We found that Ca^2+^ responses to TG2 were reduced by about 36% in neurons pre-treated with the selective of L-type blocker nifedipine (NF) (Fig. 5E-I). In addition, the drug caused a partial recovery of [Ca^2+^]_i_ towards resting levels when applied during the plateau phase of Ca^2+^ response induced by TG2 (Fig. 5E-II), suggesting that TG2 dependent Ca^2+^ influx partially occurs through L-type VOCCs.

**Figure 5.**
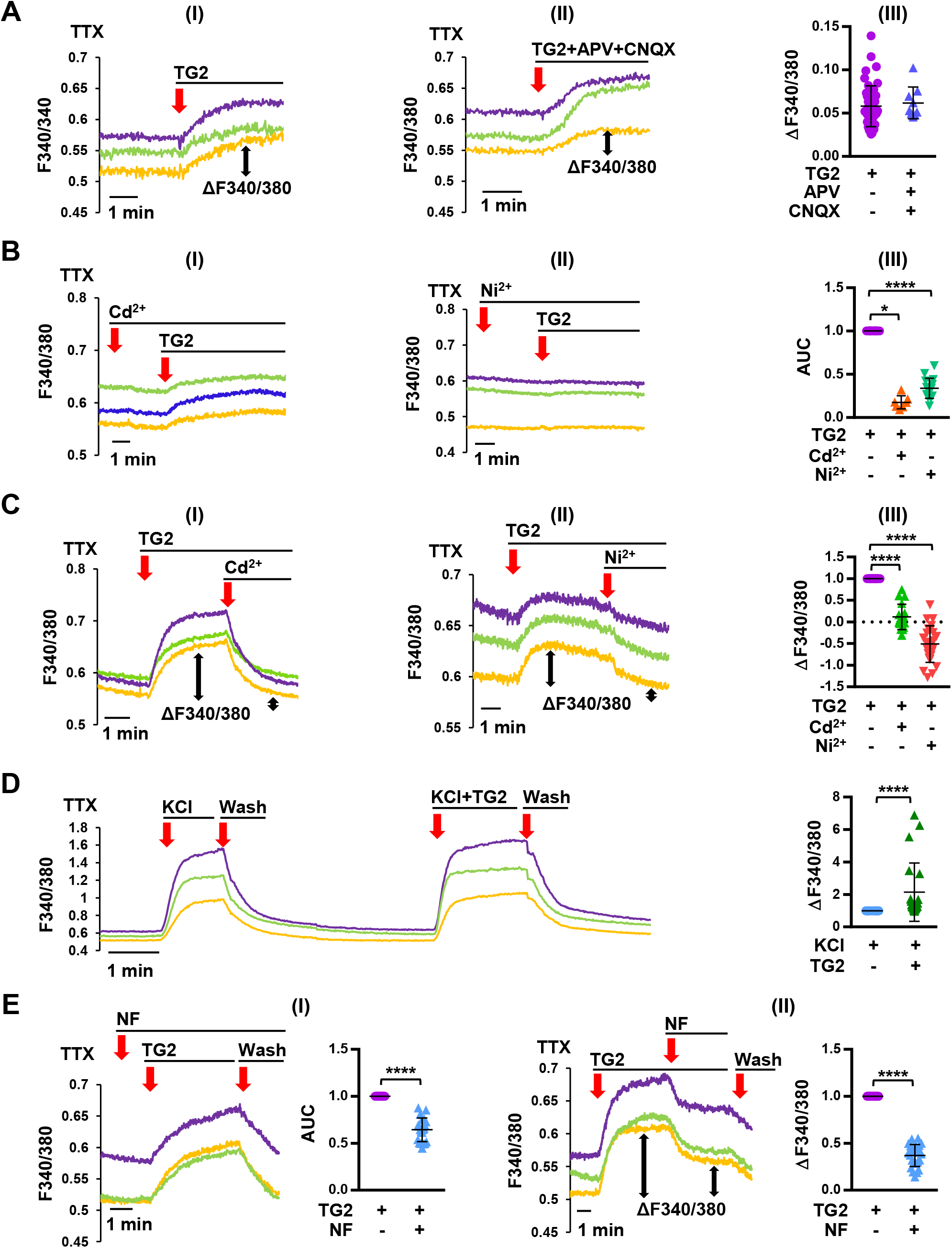
Exogenous TG2 mediates Ca^2+^ influx from the extracellular environment. **A)** Temporal analysis of [Ca^2+^]_i_ changes in 14-20 DIV neurons exposed to exogenous TG2 in the absence (I) or in the presence (II) of APV (NMDA receptor antagonist) and CNQX (AMPA/kainate receptor antagonist). TG2-dependent Ca^2+^ rise was not affected by the blockers. (III) Corresponding quantification of TG2-induced peak [Ca^2+^]_i_ rises. Data are presented as mean ΔF340/380 ± SD (N=46 and 9 neurons for TG2 alone and TG2+blockers respectively; Mann-Whitney test: p=NS). **B)** Pre-treatment with cadmium (I), a general blocker of Ca^2+^ channels, and nickel (II), a blocker of T-type VOCCs, reduced TG2-dependent Ca^2+^ responses of about 82.4% and 66.2% respectively, as quantified in (III), by measuring total Ca^2+^ influxes. Data are expressed as mean area under the curve (AUC) ± SD of TG2 + blockers normalised to TG2 alone (N=7 and 15 neurons for treatment with Cd^2+^ and Ni^2+^ respectively Wilcoxon signed-rank test: *p<0.05; ****p<0.0001). **C)** Addition of cadmium (I) and nickel (II) during the plateau phase of TG2-induced Ca^2+^ response led to a recovery of [Ca^2+^]_i_ towards basal levels. (III) Histograms show residual mean Ca^2+^ responses after blockers normalised to peak Ca^2+^ response evoked by TG2 ± SD (N=18 and 25 neurons for treatment with Cd^2+^ and Ni^2+^ respectively; Wilcoxon signed-rank test: ****p<0.0001). **D)** Ca^2+^ transients induced by 15 mM KCl in the presence of NMDAR and ampa/kainate receptor blockers APV and CNQX revealed that [Ca^2+^]_i_ rose faster and led to a 2-fold higher total Ca^2+^ influx in the presence of TG2. Data are expressed as mean AUC of TG2 + KCl normalised to KCl alone treatment ± SD (N=22 neurons; Wilcoxon signed-rank test: ****p<0.0001). **E)** Temporal analysis of [Ca^2+^]_i_ changes induced by TG2 in the presence of L-type VOCCs blocker Nifedipine (NF). (I) Total Ca^2+^ influxes in response to TG2 were reduced of about 36% in neurons pre-treated with NF. Data are expressed as mean AUC ± SD induced by TG2 in the presence of NF normalised to TG2 alone (N=21 neurons; Wilcoxon signed-rank test: ****p<0.0001). (II) NF caused a partial recovery of about 63% of Ca^2+^ concentration towards resting levels when applied during the plateau phase of Ca^2+^ response induced by exogenous TG2. Histograms show mean residual Ca^2+^ responses ± SD after NF normalised to peak Ca^2+^ response evoked by TG2 alone (N=25 neurons; Wilcoxon signed-rank test: ****p<0.0001).

### Extracellular TG2 induces membrane depolarisation and the generation of ionic inward currents, which are inhibited by Nifedipine

To further characterise the [Ca^2+^]_i_ response evoked by TG2, we performed whole cell electrophysiological recordings of hippocampal neurons. In current clamp, TG2 perfusion induced a slow membrane depolarisation (of about 20 mV) (Fig. 6A-I), which was reverted by addition of nickel (Fig. 6A-II). In voltage clamp, TG2 promoted excitatory postsynaptic currents (EPSCs), consistent with the activation of an inward Ca^2+^ current (Fig. 6B). In the same configuration, analysis of the current/voltage relationship (I/V curves) using a protocol for isolation of L-type VOCCs, showed that perfusion of TG2 (Fig. 6C-I, red I/V curve) led to an increased inward current compared to control (black I/V curve), which was completely prevented by the presence of the L-type VOCCs blocker NF (Fig. 6C-II, red I/V curve compared to control black I/V curve). These data revealed L-type VOCCs as one of the possible main targets of TG2 responsible for dysregulation of basal Ca^2+^ concentration in neurons.

**Figure 6.**
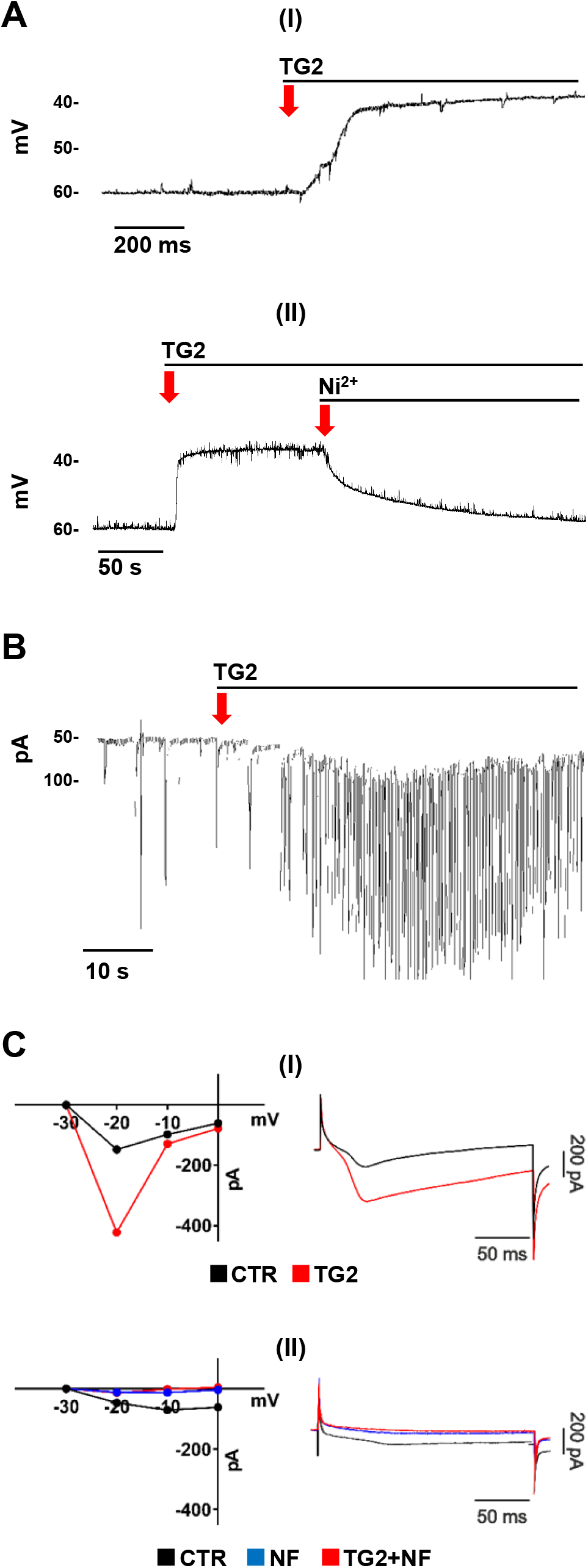
TG2 activates an inward calcium current through L-type VOCC membrane depolarisation. **A)** Whole cell current clamp. (I) Addition of TG2 induced a slow membrane depolarisation of about 20 mV (N=3). (II) Addition of Nickel after TG2 re-established resting membrane potential. **B)** Voltage clamp. TG2 perfusion promoted excitatory postsynaptic currents (EPSCs) consistent with the activation of an inward Ca^2+^ current (N=3). **C)** Whole cell voltage clamp results expressed as current/voltage (I/V) curves. (I) Perfusion of TG2 (red I/V curve) led to an increased inward current compared to control (black I/V curve). The graph on the right shows current recordings of the same experiment at −20 mV. The protocol for isolation of L-type VOCCs was applied as described in the Methods. (II) In the presence of NF (blue I/V curve), TG2 perfusion (red I/V curve, left panel) did not increase inward current compared to control (black I/V curve). The graph on the right shows current recordings of the same experiment at −20 mV (N=3).

### Contribution of the sodium/calcium exchanger to the calcium response evoked by TG2

Data so far have identified L-type VOCCs as the main channel through which extracellular TG2 induces Ca^2+^ influx and membrane depolarisation. Despite electrophysiological recordings indicated complete inhibition of inward Ca^2+^ current evoked by TG2 under block of L-type VOCCs (Fig. 6C), a residual Ca^2+^ response was observed in Ca^2+^ imaging experiments, largely sensitive to both cadmium and nickel (Fig. 5E). In addition to VOCCs, both cadmium and nickel are known to inhibit the activity of the Na^+^/Ca^2+^-exchanger (NCX), a key regulator of Ca^2+^ transport through the plasma membrane. NCX normally removes Ca^2+^ from neurons in exchange for Na^+^, which enters the neuron down its gradient across the plasma membrane (Blaustein & Lederer, 1999). However, perturbation of the Na^+^ gradient leads to operation of NCX in the reverse mode, causing Ca^2+^ influx into the neurons (Blaustein & Lederer, 1999). To investigate the involvement of NCX in the reverse mode in TG2-induced Ca^2+^ influx, we removed Na^+^ (and potassium K^+^) from the extracellular saline. In these conditions, the response to extracellular TG2 was strikingly increased (about 8-fold increase) (Fig. 7A), suggesting that NCX may amplify TG2-dependent Ca^2+^ entry. Conversely, addition of the NCX inhibitor YM-244769 significantly decreased TG2-dependent Ca^2+^ influx in both normal and Na^+^-free medium (Fig. 7B), further supporting NCX contribution to Ca^2+^ elevations in neurons exposed to TG2.

**Figure 7.**
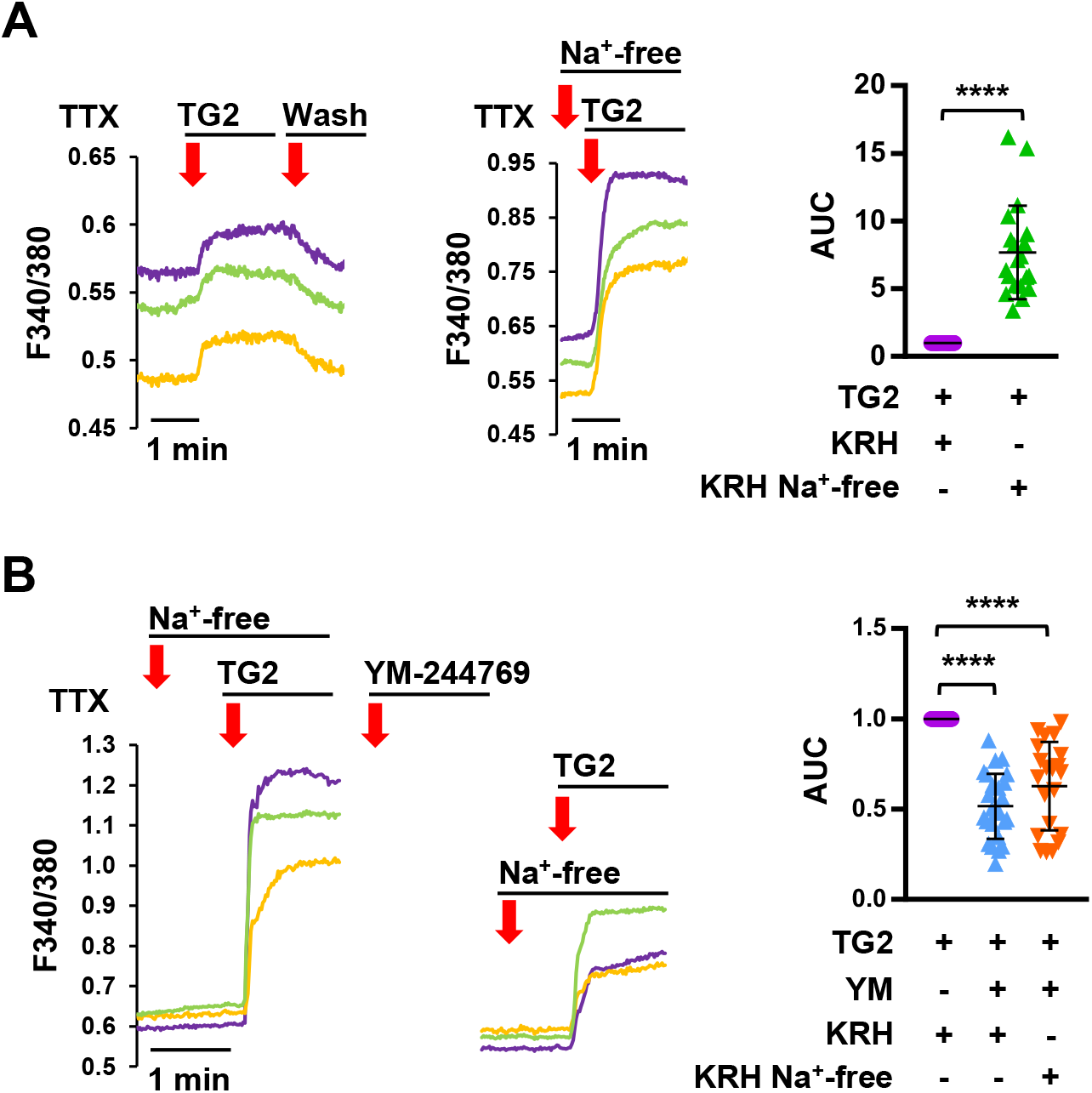
Contribution of NCX to TG2-dependent calcium rise. Temporal analysis of [Ca^2+^]_i_ changes in 14-20 DIV neurons exposed to exogenous TG2. **A)** Addition of TG2 in Na^+^-free KRH led to an 8-fold increase in average Ca^2+^ influx compared to normal KRH (N=20 neurons; Wilcoxon signed-rank test: ****p<0.0001). Data are expressed as mean ± SD normalised to TG2 AUC values. **B)** Representative temporal analysis of neuronal [Ca^2+^]_i_ responses evoked by TG2 in the absence and in the presence of the NCX inhibitor YM-244769 (10 min) in Na^+^- free KRH. YM-244769 induced a significant decrease in TG2-dependent Ca^2+^ influx. The graph shows mean TG2-induced AUC ± SD after YM-244769 normalised to TG2 alone in normal KRH (N=25 neurons; Wilcoxon signed-rank test: ****p<0.0001) and Na^+^-free KRH (N=21 neurons; Wilcoxon signed-rank test: ****p<0.0001).

### Interactors of extracellular TG2 in hippocampal neurons: Na^+^/K^+^-ATPase inhibition by extracellular TG2

To gain insights into the molecular interactions leading to the opening of L-type VOCCs and NCX operation by extracellular TG2 in neurons, we immunoprecipitated TG2 from the lysate and secretome of neurons treated with extracellular TG2 and analysed the immunoprecipitated proteins by quantitative comparative proteomics relative to untreated neurons immunoprecipitates as shown in Fig. 8A (full list of proteins is available in Table EV1).

**Figure 8.**
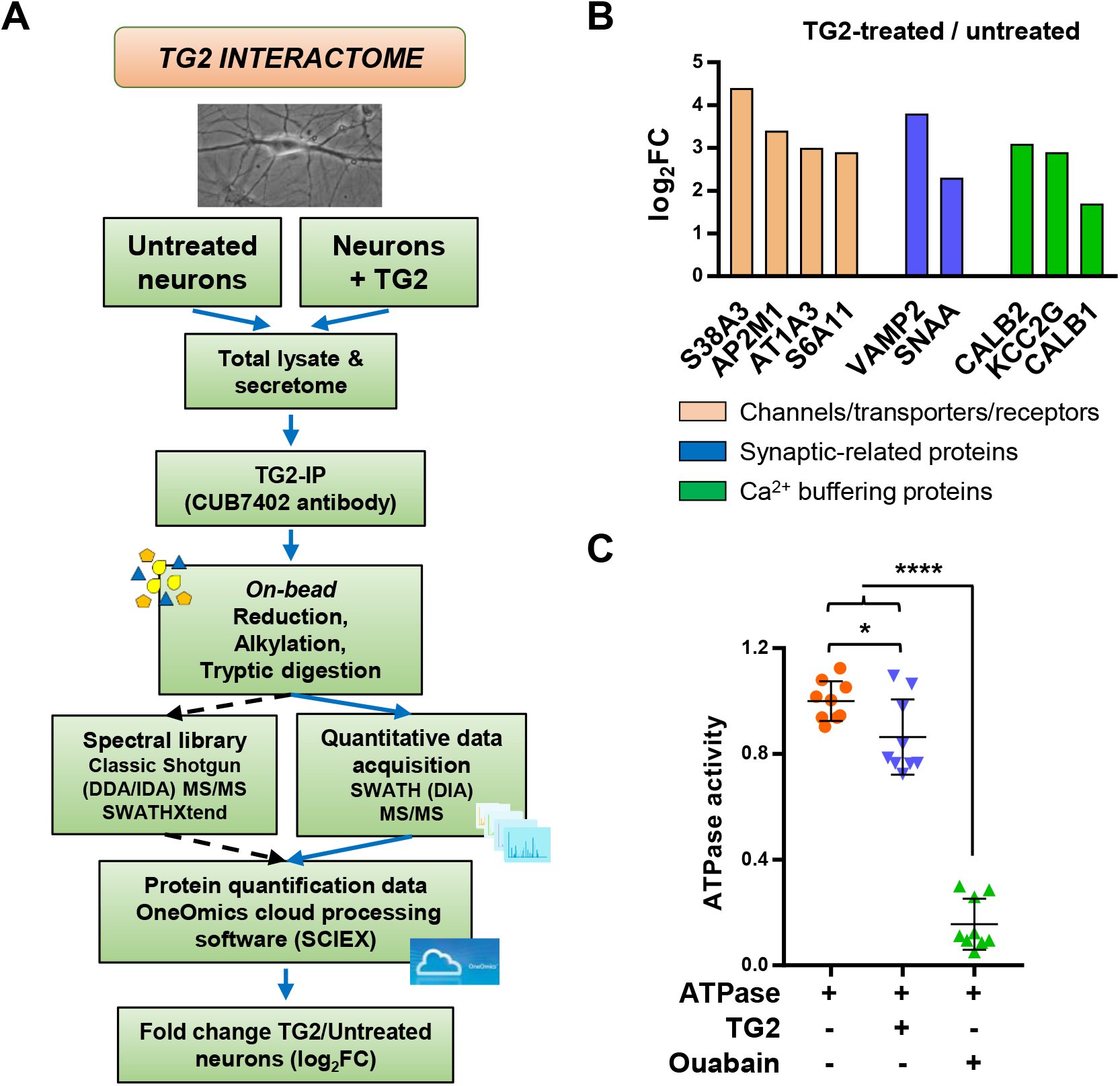
TG2 interactors in the neuronal extracellular environment: inhibition of Na^+^/K^+^-ATPase. **A)** Workflow of the identification of TG2-interactors. TG2 was immunoprecipitated from hippocampal neurons (total lysate including the secretome) incubated with extracellular TG2 or untreated (N=3 independent immunoprecipitations, from a neuronal preparation derived from ≥10 rat E18 embryos). TG2 co-precipitated proteins (TG2-IP) were trypsin digested on beads and analysed by SWATH MS. SWATH quantitative data were extracted using a spectral library produced by shotgun/data dependent acquisition (DDA) MS on all TG2-IP samples. **B)** Selection of TG2 interactors shown according to logarithm of fold change (log_2_FC) between neurons exposed to extracellular TG2 and untreated cells. **C)** The activity of porcine cerebral cortex ATPase (0.5 mU/well) assayed in the absence or presence of either gplTG2 (0.5 mU/well, 43.9 nM) or Ouabain (1 mM). Data are shown as mean ± SD of ATPase activity (mU/ml), normalised to ATPase alone values (N=3 independent experiments, one-way ANOVA test: *p<0.05; ****p<0.0001).

Among the interactors of extracellular TG2 (Fig. 8B) the finding of vesicle-associated membrane protein 2 (VAMP-2) and alpha-soluble NSF attachment protein (SNAA) (confidence >70%), suggests that extracellular TG2 localises at synapses similarly to the endogenous protein (Fig. 1). The association of Ca^2+^ buffer proteins calretinin, calbindin and a subunit of Ca^2+^/calmodulin-dependent kinase with extracellular TG2 is consistent with the ability of extracellular TG2 to elicit [Ca^2+^]_i_ changes in neurons. Among the transporters which emerged as significant TG2 partners, Na^+^/K^+^-transporting ATPase subunit α (AT1A3) was three times more associated with neurons exposed to extracellular TG2 (confidence 70%) (Fig. 8B). We hypothesised that AT1A3 could be part of the mechanism of Ca^2+^ influx triggered by extracellular TG2 via inhibition of Na^+^/K^+^-ATPase by TG2, with increase of intracellular Na^+^ leading to switch of NCX in the reverse mode (Fig 7A-B). To test ATPase inhibition by TG2, ATPase activity was measured using ATPase isolated from porcine cerebral cortex either in the presence or absence of TG2 using the specific Na^+^/K^+^-ATPase inhibitor ouabain as a control. Incubation with TG2 led to a small but significant reduction in ATPase activity suggesting that perturbation of Na^+^/K^+^-transport by extracellular TG2 may trigger the rise in neuronal Ca^2+^ influx (Fig 8C).

According to our findings, reactive astrocytes control intraneuronal [Ca^2+^]_i_ through release of TG2 in association to extracellular vesicles (Fig. 9). Upon interaction with neurons extracellular TG2 promotes the opening of L-type calcium channels and induces the Na^+^/Ca^2+^-exchanger to operate in the reverse mode, likely through inhibition of Na^+^/K^+^-ATPase, setting basal [Ca^2+^]_i_ at higher levels and enhancing neurotransmission. These findings link astrogliosis, which typically occurs under brain inflammation and degeneration with Ca^2+^ dyshomeostasis in neurons and identifies extracellular TG2 as a key astrocyte-derived factor driving neuron dysfunction.

**Figure 9.**
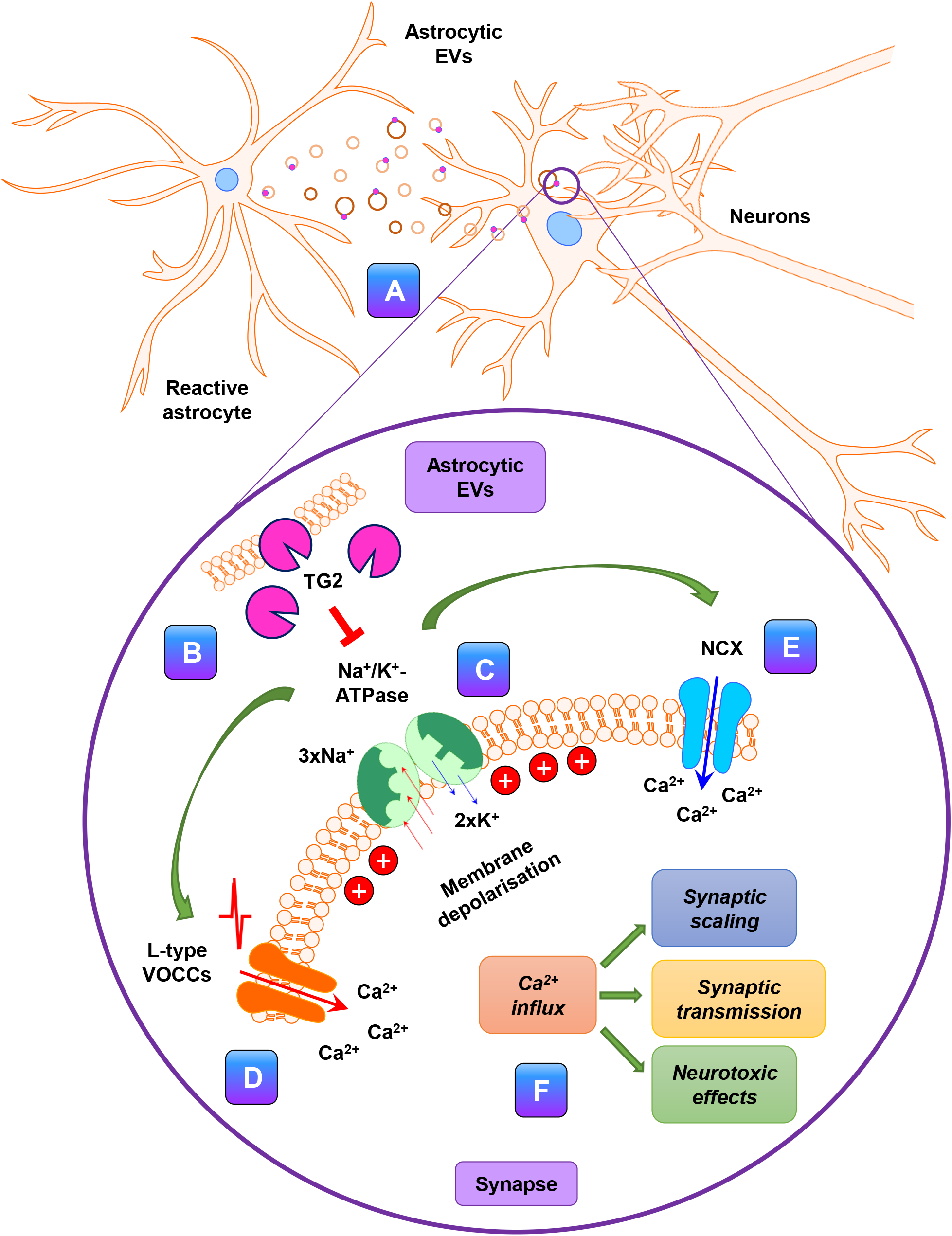
Schematic representation of the proposed mechanism. **A)** Activated astrocytes release EVs containing TG2 as a cargo and displayed at the EVs surface. **B)** Extracellular TG2 (either astrocytes vesicle-bound or free in the medium) interacts with neurons and increases [Ca^2+^]_i_ via **(C)** inhibition of Na^+^/K^+^-ATPase, causing membrane depolarisation (about 20 mV) and activation of an inward Ca^2+^ current through **(D)** L-type VOCCs and **(E)** the Na^+^/Ca^2+^ exchanger (NCX) leading to **(F)** increase of [Ca^2+^]_i_ and Ca^2+^ dyshomeostasis.

## Discussion

Our study unveils a novel mechanism by which TG2 regulates Ca^2+^ homeostasis in hippocampal neurons. We show that extracellular TG2 drives Ca^2+^ influx into neurons both in culture and brain slice, via inhibition of Na^+^/K^+^-ATPase activity and consequent activation of L-type VOCCS and NCX operating in the reverse mode.

A key observation is that the sources of extracellular TG2 are astrocytes-derived EVs, which have TG2 as cargo displayed at the EVs surface. EVs-associated TG2 significantly increases [Ca^2+^]_i_ in neurons, as demonstrated by selective [Ca^2+^]_i_ rises in response to EVs derived from WT but not TG2KO cells. Notably, TG2 content tends to increase in EVs released by reactive astrocytes. This is consistent with published data showing higher externalisation of astrocytic TG2 in response to neuroinflammation (i.e. TNF-α + IL-1β treatment) (Pinzon et al., 2017) and increased TG2 levels in association with astrogliosis, an inflammatory processes which is activated in brain in response to injury and the release of pro-inflammatory cytokines (Hostenbach, Cambron et al., 2014; Ientile et al., 2015). Astrocytes play a key role in synaptic scaling, i.e. the adjustment of synaptic strength in response to prolonged changes in the electrical activity to maintain neuronal function and survival (Papouin, Dunphy et al., 2017). Potentiation of synaptic strength largely occurs via insertion of postsynaptic AMPA receptors driven by the release of TNFα by glial cells (Stellwagen & Malenka, 2006). However, the exact mechanism and molecular actors involved in synaptic scaling are still unknown (Papouin et al., 2017). By enhancing neuronal [Ca^2+^]_i_, TG2 released in association with astrocytic EVs may favour excitatory transmission, thereby contributing to synaptic scale up. Further work is required to address this intriguing hypothesis.

EVs play an emerging role in neuron-glia cross-talk (Holm, Kaiser et al., 2018; Turola, Furlan et al., 2012). In particular, astrocyte-derived EVs induce a long lasting potentiation of spontaneous excitatory transmission (Antonucci, Turola et al., 2012; Gabrielli, Battista et al., 2015; You, Borgmann et al., 2020) but upon chronic exposure EVs from reactive astrocytes exert detrimental effects on synapse stability (Prada, Gabrielli et al., 2018), neurite differentiation and neuronal firing (You et al., 2020). Astrocyte-derived EVs isolated from plasma of Alzheimer’s disease (AD) patients carry high levels of complement effector proteins compared to healthy controls, suggesting that they could promote the spreading of inflammatory signals (Goetzl, Schwartz et al., 2018). By showing that TG2 sorted on the surface of astrocytic EVs has the capacity to regulate neuronal Ca^2+^ homeostasis, we add an important piece of information to decipher the complex astrocyte-to-neuron signalling mediated by EVs under physiology and neuroinflammation.

The action of TG2 displayed in astrocytes-derived EVs is also reproduced by soluble extracellular TG2. TG2-dependent Ca^2+^ influx sets basal [Ca^2+^]_i_ at higher levels and promotes spontaneous EPSCs. This is a novel function that has never been attributed to TG2 before. Interestingly, [Ca^2+^]_i_ is restored to basal levels upon removal of TG2 from the extra-neuronal solution, revealing that the process is reversible and not linked to neuron damage. Reversibility of TG2 action suggests also that conformation rather than transamidating activity of the enzyme regulates [Ca^2+^]_i_ homeostasis, as covalent crosslinking of a substrate would lead to irreversible effects. Several past and present evidences suggest that TG2 is capable of crosslinking-independent modifications (Akimov, Krylov et al., 2000; Stephens, Grenard et al., 2004; Steppan, Bergman et al., 2017; Verderio et al., 2003) and TG2 conformation is important in mediating the cellular response (Katt, Antonyak et al., 2018). In mouse striatal cells, TG2 in open conformation was shown to have cytotoxic effects, increasing the cells susceptibility to oxygen/glucose deprivation-mediated cell death. However, the underlying mechanism remains largely unknown (Colak, Keillor et al., 2011; Singh, Zhang et al., 2016).

Consistent with a crosslinking-independent role of TG2 in the control of Ca^2+^ dynamics, *in vitro* measurement of purified TG2 activity in KRH, mimicking the extracellular solution used for Ca^2+^ and electrophysiological recordings, showed that the protein is mostly inactive under this experimental condition. Moreover, a specific extracellular TG2 inhibitor (BOCDON) does not affect extracellular TG2-mediated [Ca^2+^]_i_ changes. On the other hand, we cannot exclude that the enzyme transamidating activity may contribute, in part, to the control of neuronal Ca^2+^ homeostasis. For example, it could be hypothesised that residual TG2 activity may act on substrates present in the extracellular medium that could promote Ca^2+^ influx upon transamidation and which would be removed together with TG2 during the washing step, thus resulting in reversibility. In this respect, transamidation of monoamine neurotransmitters, a reaction called monoaminylation, is a less characterised function of TG2 that may play a role in the process (Muma, 2018). Therefore, further work is required to exclude the possible contribution of TG2 activity to the control of Ca^2+^ dynamics.

Importantly, we show that in physiological solution TG2 induces a small increase in neuronal [Ca^2+^]_i_, not associated to cell death, setting [Ca^2+^]_i_ to levels typically displayed by astrocytes in primary culture, that are significantly higher compared to neurons. Such increase in [Ca^2+^]_i_, is functionally relevant as it promotes neuronal transmission (spontaneous EPSCs). This suggests that TG2 may exert a fundamental homeostatic function in neurons, promoting neurotransmission, not only contributing to neurodegenerative and neuroinflammatory processes upon Ca^2+^ overload (Grosso & Mouradian, 2012).

Identification of L-type VOCCs and NCX as the TG2 molecular targets mediating Ca^2+^ influx and setting basal [Ca^2+^]_i_ at a higher levels in neurons, provides mechanistic understanding of the pathway. Both molecular targets have been identified using pharmacological blockers combined to Ca^2+^ and electrophysiological recordings. Interestingly, activation of L-type VOCCs is important for synaptic plasticity (Grover & Teyler, 1990; Raymond & Redman, 2002) such as long-term potentiation (LTP), suggesting that through this target TG2 may have a role in physiological brain functions. However, alterations in VOCCs activity have been also reported in neurodegeneration, such as in AD (LaFerla, 2002; Nimmrich & Eckert, 2013), suggesting that persistent activation of VOCCs may also contribute to degenerative processes. Similarly to VOCCs, NCX is fundamental for the regulation of [Ca^2+^]_i_ and activation of the reverse mode (Ca^2+^ influx) has been associated with neuropathic pain caused by nerve-injury (Jaggi & Singh, 2011), ischemic damage (Matsuda, Arakawa et al., 2001), increased oxygen/glucose deprivation-mediated neuronal damage and infarct injury (Secondo, Pignataro et al., 2015). Thus, sustained elevations of TG2 in the extracellular environment, via opening of L-type VOCCs and NCX reverse operation, could participate to general dysregulation of Ca^2+^ homeostasis in neurodegenerative conditions. Analysis of the TG2 interactome has identified a subunit of Na^+^/K^+^-transporting ATPase as partner of extracellular TG2, and our data suggest that its inhibition by TG2 could reverse the activity of NCX, thus leading to Ca^2+^ inward flux and the opening of L-type VOCCs.

In conclusion, our data highlight for the first time a specific function of extracellular TG2, either released by astrocytes via EVs or in soluble form, on Ca^2+^ homeostasis in neurons, through opening of L-type VOCCs and NCX operation in the reverse mode, likely mediated by the Na^+^/K^+^-transporting ATPase (Fig. 8). This is the first evidence that astrocytes control intraneuronal Ca^2+^ through release of TG2 in association with EVs.

## Materials and Methods

### Primary cultures

primary cultures of hippocampal neurons were established from E18 Sprague Dawley rat embryos as previously described (Gabrielli et al., 2015). After 14-21 days in vitro (DIV), the cultures contain a significant percentage of astrocytes (Verderio, Bacci et al., 1999). Primary astrocytes from C57BL/6 WT and TG2 knockout (TG2^-/-^ or TG2KO) P3 mouse pups were established as previously described (Gabrielli et al., 2015). TG2KO mice were originally obtained from Gerry Melino (De Laurenzi & Melino, 2001) and fully backcrossed as previously described (Scarpellini, Huang et al., 2014). For TG2 cell-surface activity assay, primary astrocytes were obtained from BrainBits (BrainBits, Springfield, IL, USA) and processed following the manufacturer’s protocol.

### Reagents

guinea pig liver TG2 (Zedira GmbH, Darmstadt, Germany) 10-30 μg/ml; LPS (Sigma-Aldrich, St. Louis, MO, USA) 1 µg/ml; TTX (Tocris, Bristol, United Kingdom) 1 µM; BOCDON (Zedira) 200 µM; Nifedipine (Sigma-Aldrich) 1 µM; Cadmium (Sigma-Aldrich) 200 µM; Nickel (Sigma-Aldrich) 100 µM; YM-244769 (Tocris) 1 µM; APV (Tocris) 50 µM; CNQX (Tocris) 10 µM; biotin-cadaverine (Sigma-Aldrich) 0.1 mM; apyrase (Sigma-Aldrich) 30 U/ml; human fibronectin (Sigma-Aldrich) 5 µg/ml; ATPase from porcine cerebral cortex (Sigma-Aldrich) 0.5 mU; Ouabain (Tocris) 1 mM.

### Immunocytochemical staining

Cells were fixed in 4% paraformaldehyde - 4 % sucrose (w/v) and immunofluorescence staining was performed using the following antibodies: mouse monoclonal anti-TG2 (IA12 – Tim Johnson, University of Sheffield (Scarpellini et al., 2014)), guinea pig anti-VGLUT1 (Synaptic System, Goettingen, Germany), rabbit anti-GFAP (Dako, Agilent, Santa Clara, CA, USA), rabbit anti-Shank2 (Synaptic System), rabbit anti-Fibronectin (Sigma-Aldrich), rabbit anti-β-tubulin (Sigma-Aldrich) and rabbit anti-NR2B (Alomone, Jerusalem, Israel). Secondary antibodies were conjugated with Alexa-488, Alexa-555 and Alexa-633 fluorophores (Invitrogen). Further details are available in the Appendix Table S1. Coverslips were visualised by laser scanning Leica SP5 confocal microscope using 63X oil immersion objective. Successive serial optical sections (0.5 µm) were recorded over 5 µm planes and maximum projections were selected for quantification. Fluorescence intensity and co-localisation were estimated using the Leica LAS AF Lite software or ImageJ software as indicated in the Appendix Extended methods.

### Crude synaptosomes preparation

Two adult mouse cortices were homogenised in 10 ml of cold SHE buffer (4 mM HEPES pH 7.3, 320 mM sucrose, 1 mM EGTA, protease inhibitors), using a glass-Teflon homogeniser on ice. Brain homogenates were then centrifuged 1 000xg for 10 min at 4°C to obtain the Low speed supernatant (LSS). A part of LSS was saved for analysis, while the remaining volume was centrifuged at 12 500xg for 20 min at 4°C to obtain the crude synaptosomes (pellet). Pelleted synaptosomes were washed once in SHE (10 ml/brain) and centrifuged again at 12 500xg for 20 min at 4°C. The final pellet (crude synaptosomal fraction) was resuspended in 1 ml SHE buffer and quantified by BCA (Sigma-Aldrich).

### Western blotting

Samples were electrophoresed on 10% SDS polyacrylamide gels according to standard procedures. The following antibodies were used: mouse monoclonal anti-TG2 (IA12), anti-FLOT-2 (BD Biosciences, Wokingham, United Kingdom), anti-PSD-95 (Neuromab, Davis, C, USA), anti-TOM20 (Santa Cruz, Dallas, Texas, USA) and polyclonal rabbit anti-TG2 (ab421) (Abcam), anti-β-tubulin (Abcam), anti-ALIX (Covalab, Bron, France), anti-NR2B (Alomone, Jerusalem, Israel), anti-VGAT (Synaptic systems, Goettingen, Germany), anti-VGLUT1 (Synaptic systems), anti-GFAP (GeneTex). Further details are available in Appendix Table S1. Blots were developed using enhanced chemiluminescence (SuperSignal West Femto Maximum Sensitivity Substrate, Thermo Fisher Scientific) and protein band intensity was quantified by Aida Image Analyser v.4.03 (Raytest, Germany).

### Isolation of extracellular vesicles (EVs)

EVs were isolated from conditioned medium (CM) of astrocytes by differential centrifugation as previously described (Furini et al., 2018; Prada et al., 2018). The 10 000xg centrifugation was omitted to pellet large and small EVs together. Each astrocytes preparation derived from ≥10 mouse P3 pups. Specifically, 80% confluent monolayers were washed twice with PBS or KRH and incubated in serum-free media in the presence or absence of lipopolysaccharide (LPS, 1 µg/ml), for 24 h. CM was collected and supplemented with protease inhibitors before centrifugation. EVs were resuspended either intact in particle-free PBS (for Nanoparticle Tracking Analysis, TEM and TG2 activity assay) or KRH (for Ca^2+^ imaging) or lysed in RIPA buffer (for western blotting) or mild lysis buffer (0.25 M sucrose, 2 mM EDTA, 5 mM Tris-HCl, pH 7.4) (for TG2 activity assay).

### Nanoparticle tracking analysis

Nanoparticle tracking analysis (NTA) was performed using ZetaView PMX 120 (Particle Metrix, Meerbusch, Germany) and its corresponding software (ZetaView 8.04.02) to examine EVs size distribution and concentration. EVs samples, resuspended in particle-free PBS, were diluted to reach 50-200 particles/frame and analysed with a flow cell sensitivity of 80% across two cycles of 11 positions/cycle.

### Transmission Electron Microscopy

EVs isolated from untreated WT astrocytes were analysed by Transmission Electron Microscopy (TEM) as previously described (Charrin, Palmulli et al., 2020), with a few modifications. Briefly, EVs were deposited for 20 min on formvar/coated 200 mesh nickel EM grids (TAAB Laboratories Equipment Ltd, Berks, United Kingdom). Then grids were fixed with 2% paraformaldehyde/phosphate buffer pH 7.4 and processed for negative staining or single immunogold labelling. EVs were stained with mouse monoclonal anti-CD63 (Abcam) or anti-TG2 antibody (IA12) followed by 6 nm colloidal gold anti-mouse IgG (Jackson Immunoresearch Europe, Ely, United Kingdom). The negative staining was performed using a mixture of uranyl acetate/methylcellulose as previously described (Slot & Geuze, 2007). Brightfield transmission electron micrographs of the stained EVs were taken at an indicated magnification of 40 000x using a JEM-2100Plus (JEOL, Herts, United Kingdom) TEM operating at an accelerating voltage of 80 kV, digital micrographs recorded using a Rio 16 CMOS camera using GMS 3.0 (Gatan, Pleasanton, CA, USA).

### Total TG2 activity assay

TG2 activity of EVs either intact (resuspended in PBS) or lysed (resuspended in mild lysis buffer) was measured through incorporation of biotin cadaverine (BTC, Sigma-Aldrich) into FN as previously described (Jones, Nicholas et al., 1997). Samples were loaded in duplicates (N=3 independent experiments). TG2 activity was calculated by removing the absorbance values obtained from the respective buffer (either PBS or lysis buffer) and expressed as mU/µg of protein.

### Cytoplasmic calcium imaging

Intracellular Ca^2+^ levels were assessed in hippocampal neurons loaded with Fura-2/AM dye (Merck, Darmstadt, Germany) as previously described (Joshi, Turola et al., 2014) and explained in the Appendix Extended methods. Neurons were distinguished from astrocytes by morphology and response to KCl stimuli (Bacci et al., 1999). Experiments were performed in Krebs–Ringer’s HEPES solution (KRH) (125 mM NaCl, 5 mM KCl, 1.2 mM MgSO_4_, 1.2 mM KH_2_PO_4_, 2 mM CaCl_2_, 6 mM d-glucose, and 25 mM HEPES/NaOH, pH 7.4), whereas in some cases Na^+^/K^+^-free KRH was used instead, where both NaCl and KCl were isosmotically substituted for choline chloride. Ca^2+^ concentration was expressed as F340/380 fluorescence ratio as explained in the Appendix Extended methods. For each experiment, N indicates the total number of analysed neurons, either from multiple slides of the same cell preparation or from different preparations. Each neuronal preparation derived from ≥10 rat E18 embryos.

### Calcium imaging in brain slices

Slices were obtained from 10 days old CD1 mice and prepared as explained in the Appendix Extended methods. Acquisition protocols consisted of 1.2 fps time-lapse sequences of Fura-2 fluorescence with 30 ms exposure time. ROI over the field of view were selected, and the mean pixel intensity at each frame was measured. After 3 min baseline recording, 30 μg/ml TG2 (Zedira) in ACSF was perfused until plateau of the response was reached, then washed out. Analysis was performed using Metafluor software (Molecular Devices, San Jose, CA, USA).

### Transient transfection

8 DIV (days in vitro) neurons were transfected with 1.5 µg of pEGFP-N1 vector (Clontech Laboratories, Takara Bio Inc., Mountain View, CA, USA) or pEGFP-N1-TG2 vector (Furini et al., 2018) using 6 μl of Lipofectamine2000 (Thermo Fisher Scientific) in a total volume of 100 μl per slide (about 1.7×10^5^ neurons). After 45 min incubation, transfected neurons were washed with Neurobasal medium and incubated in filtered conditioned neuronal medium for 48 h before analysis.

### Whole cell patch clamp

Patch clamp experiments were performed in current clamp and voltage clamp configurations as explained in the Appendix Extended methods. For current clamp experiments, the effect of TG2 (30 μg/ml) on neuron resting potential was examined. Voltage clamp experiments were performed to observe the involvement of Voltage Operated Calcium Channels (VOCCs) in the phenomenon. Specifically, L-type VOCCs were isolated with a pre-pulse of 40 mV and a Δ10 mV step protocol from −30 to 0 mV.

### Isolation of TG2 complexes and Data Acquisition by Mass Spectrometry

TG2-associated proteins were isolated from hippocampal neurons incubated with 30 μg/ml TG2 in KRH buffer for 5 min and from untreated neurons as control (N=3). After incubation, the supernatant was removed and neurons were lysed using IP buffer (25 mM Tris-HCl, 150 mM NaCl, 1 mM EDTA, 1% [v/v] NP40 detergent solution, and 5% [v/v] glycerol, pH 7.4) containing protease inhibitor cocktail (Sigma-Aldrich). The whole cell lysate was centrifuged at 13 000xg for 10 min at 4°C to remove larger particulates. TG2 with associated proteins was immunoprecipitated from cell lysates and supernatants as previously described (Furini et al., 2018) with few modifications. Incubation of samples with the anti-TG2 antibody-coated beads (CUB7402; Thermo Fisher Scientific) was performed for 20 h at 4°C in constant rotation. Proteins were reduced, alkylated and trypsin digested directly on the beads and peptides were analysed by RP-HPLC-ESI-MS/MS using a TripleTOF 6600 mass spectrometer (SCIEX) in data-dependent acquisition mode for spectral library construction, and in SWATH 2.0 data-independent acquisition mode using 100 variable SWATH acquisition windows for quantification as described in the Appendix Extended methods. The mass spectrometry proteomics data have been deposited to the ProteomeXchange Consortium via the PRIDE (Perez-Riverol, Csordas et al., 2019) partner repository with the dataset identifier “PXD022224”. Analysis of differentially expressed proteins was performed using the OneOmics cloud processing online platform (SCIEX) as described in the Appendix Extended methods.

### ATPase activity assay

Na^+^/K^+^-ATPase activity was analysed using the High Throughput Colorimetric ATPase Assay kit (Innova Biosciences, Cambridge, United Kingdom) according to the manufacturer’s instructions. ATPase isolated from porcine cerebral cortex (0.5 U/well) was assayed in the presence or absence of gplTG2 (0.5 U/well) and specific Na^+^/K^+^-ATPase inhibitor ouabain (1 mM).

### Statistical analysis

All data are presented as mean ± SD from the indicated number of independent experiments (N=3) or cells as indicated in the respective figure legends. Each primary cell preparation was derived from ≥10 rat E18 embryos (hippocampal neurons) or ≥10 mouse P3 pups (astrocytes). Data in each experiment were first tested for normal distribution by D’Agostino-Pearson normality test (GraphPad Prism 7.05 software). In the case of normally distributed data, statistical analysis was performed by Student’s t-test (2 groups comparisons) or one-way Anova (Tukey multiple comparisons test). In the case of non-normally distributed data, statistical analysis was performed by nonparametric Mann-Whitney test (2 groups comparisons) or Kruskal-Wallis Dunn’s test (multiple comparisons). For paired normalised data, the Wilcoxon matched-pairs signed rank test was used. Differences are considered significant by p<0.05.

## Data Availability Statement

The mass spectrometry proteomics datasets have been deposited to the ProteomeXchange Consortium via the PRIDE (Perez-Riverol et al., 2019) partner repository with the dataset identifier “PXD022224” (http://www.ebi.ac.uk/pride). The rest of the data are included in this published article (and its supplementary information files).

## Acknowledgments

This work was supported by the John Turland PhD scholarship (to E.T. and E.A.M.V.), the American Association Research fellowship (to I.P. and C.V.) (AARF-588984), the IBRO/PERC InEurope Short Stay grant to E.T, the AIRC fellowship for Italy to I.V., The Marmont Foundation and the Nottingham Trent University REF-Quality Research fund (to E.A.M.V.). We are grateful to Prof Jeff Keillor (University of Ottawa, Canada) for critically reading the manuscript. We thank Prof David Abraham (University College London/Medical School, United Kingdom) and Prof Timothy Johnson (University of Sheffield, United Kingdom) for generously sharing facilities. We are grateful to Dr Giulia D’Arrigo (CNR Institute of Neuroscience, Italy) for her help in maintaining the primary cell cultures and we gratefully acknowledge the NTU Imaging Suite and Dr Graham J. Hickman for assistance in obtaining TEM data.

## Author Contribution Statement

E.T., M.M, C.V. and E.A.M.V conceived and designed research. E.T., I.V., M.G., G.F., C.C. and M.P.S performed research. E.T., I.V., I.P., M.G., D.J.B., G.S., M.M., C.V. and E.A.M.V analysed and interpreted data. E.T., I.V., C.V. and E.A.M.V. wrote the paper. All authors read and approved the final paper.

## Expanded View Figure Legends

**Figure EV1. TG2 is localised on the surface of primary astrocytes**. Cell surface TG2 activity in primary astrocytes (up to 44 DIV) using TG2 inhibitor ZDON (100 µM) to validate TG2 specific activity. Data are shown as mean ± SD of raw absorbance values minus background, normalised to CTR in the presence of 0.1% DMSO (vehicle) (N=4, Mann-Whitney test: ***p<0.001).

**Figure EV2. TG2 localises extracellularly in primary astrocytes and co-localises with fibronectin**. Immunofluorescence staining of primary astrocytes. Cells were fixed in 3% paraformaldehyde (w/v) without permeabilization and stained with anti-TG2 IA12 (green), DAPI (blue) and ECM marker fibronectin (FN, red). Coverslips were visualised by laser scanning Leica SP5 confocal microscope using 63X oil immersion objective. Successive serial optical sections (1 µm) were recorded over 8 µm planes. Scale bar 20 µm.

**Figure EV3. TG2 in KRH has negligible activity in the absence of reducing agent (DTT)**. Total TG activity assay assessing TG2 activity levels in vitro expressed as mU/µg of protein. TG2 was incubated in KRH buffer pH 7.4 (which contains 2 mM calcium) in the presence or absence of DTT 10 mM. Data is expressed as mean ± SD (N=3, each point analysed in duplicate. Mann-Whitney test: **p<0.01).

