## Appendix for "Astrocytic extracellular vesicles modulate neuronal calcium homeostasis via transglutaminase-2"

Elisabetta A.M. Verderio

### Table of content:

- Extended methods
- Supplementary Table S1

### Extended methods

***Image analysis of dual localisations:*** TG2 and VGLUT1 (or Shank-2) double-positive puncta were revealed by generating a TG2/VGLUT1 (or TG2/Shank-2) double-positive image with ImageJ. A fixed threshold was then set in the double-positive images and the number of TG2-positive puncta was measured and normalised to total number of VGLUT1 puncta or Shank-2 puncta to obtain the fraction of TG2 positive presynaptic or postsynaptic sites respectively.

***Cytoplasmic calcium imaging***: Neurons between 10 and 17 DIV were incubated with 2 µM Fura-2/AM (Merck, Darmstadt, Germany) in neuronal conditioned medium for 40 min at 37$^{o}$C, washed in Krebs–Ringer's HEPES solution (KRH) and transferred to the recording chamber of an inverted microscope (Axiovert 100, Zeiss, Oberkochen, Germany) equipped with a $Ca^{2+}$ imaging unit. Polychrome V (TILL Photonics GmbH, Graefelfing, Germany) was used as the light source with excitation at 340 and 380 nm wavelengths. Images were collected with a CCD Imago-QE camera (TILL Photonics GmbH) and analysed with FEI Live Acquisition 2.6.0.14 software. $Ca^{2+}$ concentration was expressed as F340/380 fluorescence ratio. The ratio values in selected region of interest (ROI) corresponding to neuronal somata were calculated from sequences of images to obtain temporal analysis. Peak $Ca^{2+}$ rise was quantified as ΔF340/380 while total $Ca^{2+}$ influx was measured by analysing the Area under the curve (AUC) during 30 seconds after each stimulus. In the experiment using EVs, $[Ca^{2+}]_i$ was recorded in several fields of the same slide prior to EVs addition (basal $Ca^{2+}$) and 20 min after, in order to allow EVs to deposit and interact with the neurons. Recordings were performed in the presence of APV, CNQX and apyrase to exclude the possible activation of NMDAR, AMPA/kainate receptors and ATP respectively.

***Calcium imaging in brain slices:*** CD1 mice (10 days old) were deeply anesthetized by isoflurane inhalation, kept under controlled general anaesthesia (isoflurane 2.5% − $O_2$ 70%, flow rate 2/2.5 l/min, Isoflurane Vaporizer, Ugo Basile, Gemonio, Italy) and then decapitated after tail pinch reflex disappearance. The brain was rapidly removed and 300 µm thick coronal sections containing the hippocampus were made on an OTS-5000 vibratome (EMS- Electron Microscopy Sciences,

Hatfield, PA, USA). All steps were performed in ice cold artificial CSF (ACSF; in mM: NaCl 125, $NaHCO_3$ 25, D-glucose 25, KCl 2,5, $MgCl_2$ 1, $CaCl_2$ 2, $NaH_2PO_4$ 1.25) oxygenated with 95% $O_2$/5% $CO_2$, as in (Cameron, Kekesi et al., 2016). Prior to recording, slices were stored for at least 1 h in a recovery chamber containing oxygenated ACSF solution at room temperature. After recovery, single slices were transferred to a custom-made loading chamber (3 cm diameter; 2 ml ASCF) and incubated for 45 min/1 h at 32°C with 10 μM Fura-2 AM $Ca^{2+}$ dye in ACSF bubbled with 95% $O_2$/5% $CO_2$ protected from light (adapted from (Dawitz, Kroon et al., 2011)). A single slice was then transferred to a submerged recording chamber, held under a harp-shaped slice anchor (Warner Instruments, Hamden, CT, USA) and perfused at a flow rate of 1.3 ml/min (Solution flow valve with On-Off Switch, Warner Instruments, Hamden, CT, USA) with oxygenated ACSF, at 22 ± 1°C (TC-324B automatic temperature controller, Warner Instruments, Hamden, CT, USA). Cells were visualized with an upright Nikon Eclipse FN-S2N microscope (Nikon, Tokyo, Japan) using a Nir Apo 40X Nikon objective (water-immersion, 3.5 mm working distance, 0.80 numerical aperture) (Nikon, Tokyo, Japan) and Metafluor software (Molecular Devices, San Jose, CA, USA). For ratiometric imaging of Fura-2, the excitation light was filtered through a Lambda 10-2 Wavelength Switcher (Sutter Instruments) to provide wavelengths of 340 and 380 nm. Emission light was passed through a bandpass 510 nm filter and captured by a digital camera (Photometric CoolSNAP es, Teledyne Photometrics, Tucson, AZ, USA).

***Whole cell patch clamp***: The patch electrodes (BB150F-8P with filament, Science Products) with a diameter of 1.5 mm, were pulled from hard borosilicate glass on a Brown-Flaming P-97 puller (Sutter Instrument, Novato, CA) and fire-polished to a tip diameter of 1-1.5 μm and an electrical resistance of 4-6 MΩ. Neurons were voltage-clamped using an Axopatch 200 B amplifier (Axon Instrument, Molecular Devices, San Jose, CA, USA) in the whole-cell configuration. Ionic currents were digitized at 5 kHz and filtered at 1 kHz. Clampex 9 was used as the interface acquisition program. The external solution was constituted of 140 mM NaCl, 5 mM TEA-Cl, 10 mM Hepes, 1 mM $MgCl_2$, 2 mM $CaCl_2$, 5 mM Glucose (314 mOsm). The internal solution was made of 127 mM

Cs-Gluconate, 4 mM NaCl, 10 mM Hepes, 2 mM $MgCl_2$, 0.1 mM $CaCl_2$, 10 mM Glucose, 1 mM EGTA. 4 mM Mg-ATP and 0.3 mM Na-GTP were added fresh to the solution (295 mOsm).

***Total TG2 activity assay:*** The total TG activity of gplTG2 in the presence or absence of reducing agent Dithiothreitol (DTT) was measured through incorporation of biotin cadaverine (BTC, Sigma-Aldrich) into FN as previously described (Jones, Nicholas et al., 1997). TG2 (30 µg/ml) was first diluted in KRH buffer to mimic conditions of calcium imaging experiments and then incubated 2 h at 37°C with reaction buffer containing 50 mM Tris-HCl pH 7.4, 0.1 mM BTC and either 5 mM $CaCl_2$ or 5 mM EDTA, in the presence or absence of 10 mM DTT. Samples were loaded in duplicates, to a final volume of 100 µl per well (N=3). A standard curve with known quantities of gplTG2 in reaction buffer containing DTT was also included. The reaction was stopped by addition of 10 mM EDTA in PBS, followed by 3 washes with 50 mM Tris-HCl pH 7.4. The plate was pre-incubated with 50 mM phosphate citrate buffer (Sigma-Aldrich) and the amount of BTC crosslinked by TG2 activity was revealed by addition of ExtrAvidin peroxidase, followed by 3,3',5,5'-tetramethylbenzidine (TMB, Sigma-Aldrich, St. Louis, MO, USA) and $H_2SO_4$. Spectrophotometric absorbances were measured at 450 nm. TG2 activity was calculated by removing the mean values obtained for each sample in the presence of EDTA from the mean values in the presence of $Ca^{2+}$. TG2 activity was expressed as mU/µg of protein.

***TG2 cell-surface activity assay:*** TG2 activity associated with the extracellular surface was assayed in live cells in culture by measuring the incorporation of biotin-cadaverine into fibronectin (FN), as described in (Scarpellini, Germack et al., 2009). Briefly, astrocytes were plated into 96-well plates pre-coated with human plasma FN in serum-free medium (which contains an activating concentration of $Ca^{2+}$), in the presence of the amine substrate biotinylated cadaverine (0.1 mM). Cells were allowed to adhere for 2 h at 37°C, then washed with PBS, pH 7.4, and incubated with 0.1% deoxycholate in PBS. The biotinylated cadaverine incorporated into the deoxycholate-insoluble FN matrix was revealed by incubation with ExtrAvidin peroxidase (Sigma-Aldrich) followed by addition of TMB at fixed time and stopped by 2.5 N $H_2SO_4$ (Fisher Scientific). Spectrophotometric absorbances were measured at 450 nm.

***SWATH acquisition MS and data analysis of TG2-immunoprecipitates:*** Samples (3 µl, ~ 5 µg protein digest) were directly injected by autosampler (Eksigent nanoLC 425 LC system) at 5 µL/min onto a YMC Triart-$C_{18}$ column (15 cm, 3 µm, 300 µm i.d.) using gradient elution (2–40% mobile phase B, followed by wash at 80% B and re-equilibration) over either 73 min (87 min run time) (for spectral library construction using data/information dependent acquisition DDA/ IDA) or 43 min (57 min run time) for SWATH/DIA (data independent acquisition) analysis. Mobile phases consisted of A: water containing 0.1% (v/v) formic acid; B: acetonitrile containing 0.1% (v/v) formic acid. The LC system was hyphenated to a Sciex TripleTof 6600 mass spectrometer fitted with a Duospray source and 50 µm electrode suitable for microflow proteomic analysis. The IDA method was run with parameters of: CUR 25; GS1 13; GS2 0; ISVF 5500; TEM 0. TOFMS mass range of 400–1250 *m/z*; accumulation time of 250 ms with product ion scans on the top 30 ions before switching (dynamic exclusion for 20 s) with rolling collision energy selected. Product ion accumulation time was set to 50 ms giving a cycle time of 1.8 s. The SWATH method was run with the same source parameters as the IDA with a 50 ms TOFMS scan followed by 100 variable SWATH windows (optimized on an IDA datafile of the same samples) of 25 ms between 100 and 1500 *m/z* giving a cycle time of 2.6 s.

A spectral library for SWATH data extraction was constructed using the output from ProteinPilot 5.02 (Sciex) searching against the Swissprot mouse database (January 2019), all IDA runs (pooled samples) and aligned using PeakView 2.1 (Sciex) SWATH microapp against multiple endogenous peptides. SWATH data extraction, normalization, quantitation and fold change analysis were carried out using Sciex's OneOmics cloud processing software suite incorporating processing methodology from (Lambert, Ivosev et al., 2013). Proteins were considered as TG2 specific interactors if they had quantitative data on more than a single peptide and a positive $log_2FC$ value in the TG2/Untreated comparison, with a OneOmics confidence threshold of 55%. Functional classification of the identified proteins was performed using PANTHER (Protein ANalysis THrough Evolutionary Relationships) database (www.pantherdb.org) or manually searched in UniProt (www.uniprot.org), referring specifically to the "GO - Molecular Function" annotation terms ontology.

***ATPase activity assay***: ATPase activity was assessed by the High Throughput Colorimetric ATPase activity assay (Innova Biosciences, Cambridge, United Kingdom) according to manufacturer instructions. Briefly, 0.5 mU of Triphosphatase from porcine cerebral cortex (Sigma-Aldrich) were incubated in assay buffer (50 mM Tris-HCl pH 7.4, 100 mM NaCl, 25 mM KCl, 3 mM $MgCl_2$, 2 mM $CaCl_2$, 0.5 mM ATP) either alone or in the presence of 0.5 mU of gplTG2 (Sigma-Aldrich) or 1 mM Ouabain (Tocris, Bristol, United Kingdom) for 30 min at RT. Reaction was stopped by addition of $P_i$ColorLock, followed by stabilizer and incubation at RT for further 30 min. Spectrophotometric absorbances were measured at 595 nm.

**Table S1.** Antibodies for Immunofluorescence, western blot and immunoprecipitation.

| Antibody | Manufacturer | Product code | Application | Reference / validation profile |
|---|---|---|---|---|
| Mouse monoclonal anti-TG2 (IA12 clone) | Tim Johnson, University of Sheffield | NA | IF, WB | (Scarpellini, Huang et al., 2014) |
| Mouse monoclonal anti-TG2 (Cub7402 clone) | Thermo Fisher Scientific | MA5-12739 | IP | https://www.antibodypedia.com/gene/3611/TGM2/antibody/561454/MA5-12739 |
| Rabbit polyclonal anti-TG2 | Abcam | ab421 | WB | (Sheftel & Hernandez, 2020) |
| Guinea pig polyclonal anti-VGLUT1 | Synaptic System | 135 304 | IF, WB | https://www.labome.com/product/Synaptic-Systems/135-304.html |
| Rabbit polyclonal anti-GFAP | Dako, Agilent | Z0334 | IF | (Bussian, Aziz et al., 2018) |
| Rabbit polyclonal anti-GFAP | GeneTex | GTX108711 | WB | https://www.antibodypedia.com/gene/3505/GFAP/antibody/174665/GTX108711 |
| Rabbit polyclonal anti-Shank2 | Synaptic System | 162 202 | IF | (Joshi, Turola et al., 2014) |
| Rabbit polyclonal anti-Fibronectin | Sigma-Aldrich | F3648 | IF | https://www.antibodypedia.com/gene/3522/FN1/antibody/79963/F3648 |
| Rabbit polyclonal anti-NR2B | Alomone | AGC-003 | IF, WB | https://www.labome.com/product/Alomone-Labs/AGC-003.html |
| Mouse monoclonal anti-FLOT-2 | BD Biosciences | 610383 | WB | https://www.labome.com/product/BD-Biosciences/610383.html |
| Mouse monoclonal anti-PSD-95 | Neuromab | 75-028 | WB | https://www.labome.com/knockout-validated-antibodies/PSD-95-antibody-knockout-validation-NeuroMab-75-028.html |
| Mouse monoclonal anti-TOM20 | Santa Cruz | Sc-17764 | WB | https://www.antibodypedia.com/gene/2619/TOMM20/antibody/12232/sc-17764 |
| Rabbit polyclonal anti-β-tubulin | Abcam | Ab6046 | WB | https://www.labome.com/product/Abcam/ab6046.html |
| Rabbit polyclonal anti-β-tubulin | Sigma-Aldrich | T3526 | IF, WB | (Joshi et al., 2014) |

| | | | | |
|---|---|---|---|---|
| Rabbit polyclonal anti-VGAT | Synaptic System | 131 103 | WB | https://sysy.com/product/131103 |
| Rabbit polyclonal anti-ALIX | Covalab | Pab0204 | WB | (D'Arrigo, Gabrielli et al., 2021) |
| Secondary antibodies anti-mouse Alexa-488, anti-rabbit Alexa-555 and anti guinea pig Alexa-633 | Invitrogen | A32723, A32732, A-21105 | IF | NA |

IF: Immunofluorescence; WB: western blot; IP: immunoprecipitation.
