## Supplementary figures and images for "Astrocytic extracellular vesicles modulate neuronal calcium homeostasis via transglutaminase-2"

### Fig. EV1

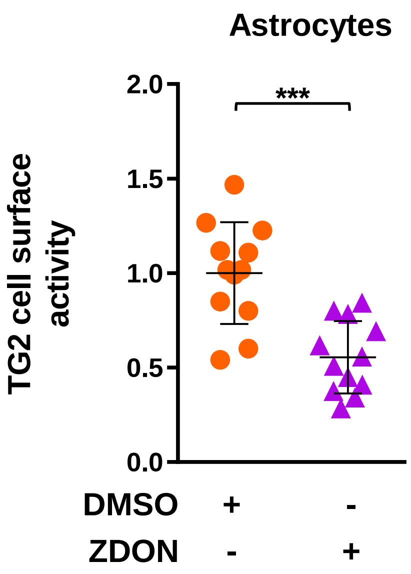

### Fig. EV2

#### Figure EV2

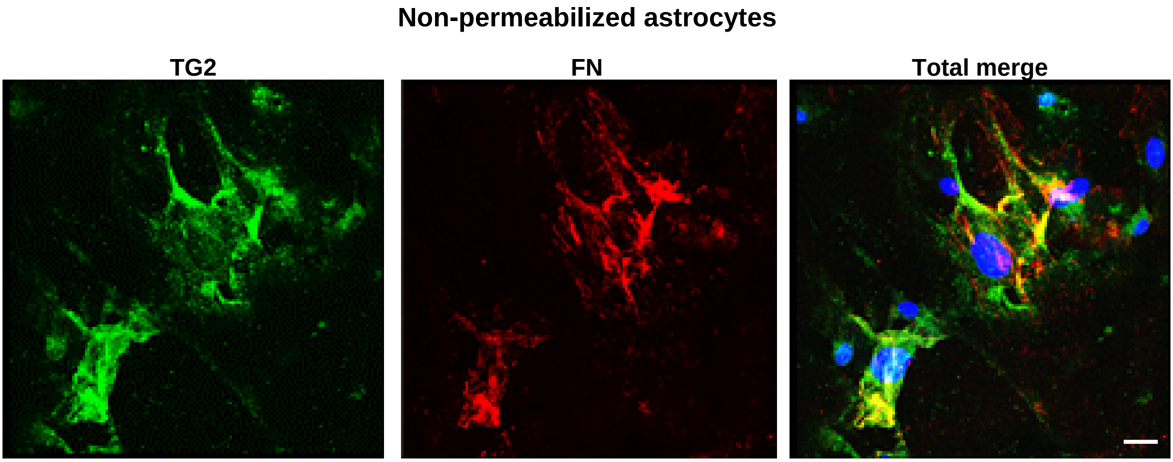

### Fig. EV3

##### Figure EV3

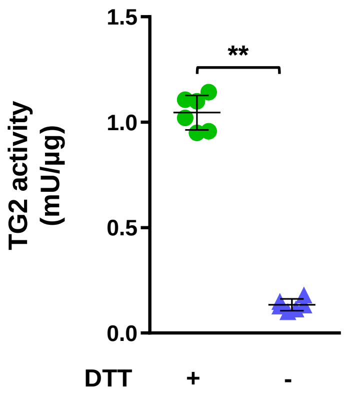
